# Co-condensation of proteins with single- and double-stranded DNA

**DOI:** 10.1101/2021.03.17.435834

**Authors:** Roman Renger, Jose A. Morin, Regis Lemaitre, Martine Ruer-Gruss, Frank Jülicher, Andreas Hermann, Stephan W. Grill

## Abstract

Biomolecular condensates provide distinct compartments that can localize and organize biochemistry inside cells. Recent evidence suggests that condensate formation is prevalent in the cell nucleus. To understand how different components of the nucleus interact during condensate formation is an important challenge. In particular, the physics of co-condensation of proteins together with nucleic acids remains elusive. Here, we use optical tweezers to study how the prototypical prion-like protein Fused-in-Sarcoma (FUS) forms liquid-like assemblies in vitro, by co-condensing together with individual DNA molecules. Through progressive DNA unpeeling, buffer exchange and force measurements, we show that FUS adsorbing in a single layer on DNA effectively generates a sticky FUS-DNA polymer that can collapse to form a liquid-like FUS-DNA co-condensate. Condensation occurs at constant DNA tension for double-stranded DNA, which is a signature of phase separation. We suggest that co-condensation mediated by protein adsorption on nucleic acids is an important mechanism for intracellular compartmentalization.

## Introduction

Many cellular compartments that provide distinct biochemical environments are not separated by a lipid membrane. An important class of such membrane-less compartments are formed by the condensation of proteins and other components in dynamic assemblies called biomolecular condensates (Hyman, Weber and Jülicher, 2014; Aguzzi and Altmeyer, 2016; Banani *et al.*, 2017). Biomolecular condensates increase the local concentration of their components, which can lead to substantially accelerated biochemical reactions (Li *et al.*, 2012; Hernández-Vega *et al.*, 2017). Condensates that form beyond a saturation concentration can buffer the cellular concentration of molecules while at the same time clamping the concentration of phase-separated components inside (Klosin *et al.*, 2020). Biomolecular condensates could also localize reaction components, and by excluding molecules from condensates they can contribute to enhance specificity of biochemical processes. The formation of biomolecular condensates often relies on the existence of low-complexity domains (Han *et al.*, 2012; Kwon *et al.*, 2013; Patel *et al.*, 2015; Wang *et al.*, 2018). Condensates can show liquid-like material properties: they deform under shear stress, fuse, round up and exchange their constituents with the environment (Brangwynne *et al.*, 2009; Jawerth *et al.*, 2018, 2020).

Many condensed structures play essential roles in nuclear organization. For example, heterochromatin is a dense form of chromatin in which DNA co-condenses with specific factors as well as nucleosomes to form transcriptionally silent domains of chromatin (Larson *et al.*, 2017; Strom *et al.*, 2017; Larson and Narlikar, 2018; Sanulli *et al.*, 2019; Keenen *et al.*, 2021). Furthermore, transcriptional hubs, or condensates, are dense and dynamic assemblies of transcription factors, associated proteins, DNA and RNA. Such condensates have been suggested to play an important role in the generation of transcriptional hubs that could coordinate the expression of several genes and mediate enhancer function (Hnisz *et al.*, 2017; Cho *et al.*, 2018; Sabari *et al.*, 2018; Guo *et al.*, 2019; Henninger *et al.*, 2021). Recently it was shown that a pioneer transcription factor can form co-condensates together with DNA *in vitro* (Quail *et al.*, 2020). Some membrane-less compartment in the cell nucleus, such as the nucleolus, show all the features of liquid-like condensates (Brangwynne, Mitchison and Hyman, 2011; Feric *et al.*, 2016). However, for the majority of smaller nuclear compartments, the physical mechanisms by which they form remain controversial. In particular, the physicochemical mechanisms that drive co-condensation of proteins together with nucleic acids remain not well understood.

A prominent nuclear condensate is formed after DNA damage, where multiple proteins come together at the damage site to repair DNA (Aleksandrov *et al.*, 2018; Levone *et al.*, 2021). Early components of the DNA damage condensate are members of the FET family such as the prion-like protein Fused-in-Sarcoma (FUS) (Altmeyer *et al.*, 2015; Patel *et al.*, 2015; Naumann *et al.*, 2018). FUS has been shown to form liquid-like condensates in bulk solution at μM FUS concentrations (Patel *et al.*, 2015; Maharana *et al.*, 2018). However, its role in forming DNA repair compartments remains unknown.

FUS is a modular protein that consists of a nucleic acid binding domain containing various nucleic acid binding motifs and an intrinsically disordered low-complexity domain that mediates FUS self-interaction (Schwartz *et al.*, 2013; Wang, Schwartz and Cech, 2015) It is involved in a multitude of physiological intracellular processes related to nucleic acid metabolism, for example transcriptional regulation (Tan *et al.*, 2012; Yang *et al.*, 2014), mRNA splicing (Rogelj *et al.*, 2012), processing of non-coding RNA (Shelkovnikova *et al.*, 2014), DNA damage response (Aleksandrov *et al.*, 2018; Naumann *et al.*, 2018; Singatulina *et al.*, 2019; Levone *et al.*, 2021), ensuring mRNA stability (Kapeli *et al.*, 2016), mRNA trafficking (Fujii and Takumi, 2005) and regulation of mRNA translation under stress conditions (Li *et al.*, 2013). FUS also forms higher order aggregated and oligomeric assemblies in a set of neurodegenerative disorders (Patel *et al.*, 2015; Naumann *et al.*, 2018; Alberti and Dormann, 2019)

While performing its physiological tasks, FUS typically acts in dynamic assemblies that are formed with or on nucleic acids or nucleic acid-like polymers. In the context of DNA damage, the formation and dissolution of FUS condensates depends on the presence or absence of poly(ADP-ribose) (PAR), a DNA-like sugar polymer produced by PAR polymerases (Altmeyer *et al.*, 2015; Patel *et al.*, 2015; Aleksandrov *et al.*, 2018; Naumann *et al.*, 2018; Singatulina *et al.*, 2019). Other examples for FUS-enriched condensates are stress granules, which are liquid-like, dynamic cytoplasmic hubs that form upon heat stress (Li *et al.*, 2013; Patel *et al.*, 2015; Protter and Parker, 2016) or nuclear granules, which are associated with transcription and splicing (Patel *et al.*, 2015; Thompson *et al.*, 2018)

To investigate the physics underlying FUS-DNA condensate formation, we devised an *in vitro* assay based on optical tweezers combined with confocal microscopy. This allowed us to manipulate single DNA molecules in the presence of FUS protein in solution, image FUS proteins associating with the DNA molecule, and at the same time control and measure pN forces exerted on the DNA.

## Results

We set out to establish a biophysical assay based on optical tweezers and confocal microscopy to investigate collective interactions between FUS and DNA. For this, we exposed individual lambda phage DNA molecules stretched between two polystyrene beads each held in place in an optical trap to FUS-EGFP (from here on called “FUS”) inside a microfluidics flow chamber (Figure 1A). Scanning confocal fluorescence microscopy was used to visualize the binding of FUS to DNA (van Mameren *et al.*, 2009; Candelli *et al.*, 2014; Brouwer *et al.*, 2016). We first trapped two streptavidin-coated polystyrene beads, which were then used to catch and stretch a lambda phage double stranded DNA (dsDNA) molecule that was biotinylated at the two termini of only one of its two complementary strands. Next, we verified that indeed only a single DNA molecule was stretched by evaluating the mechanical properties of the connection and comparing it to the properties of a single lambda phage DNA molecule (see below). Finally, we exposed the stretched DNA molecule to bulk FUS protein while imaging the system with a scanning confocal fluorescence microscope.

**Figure 1.**
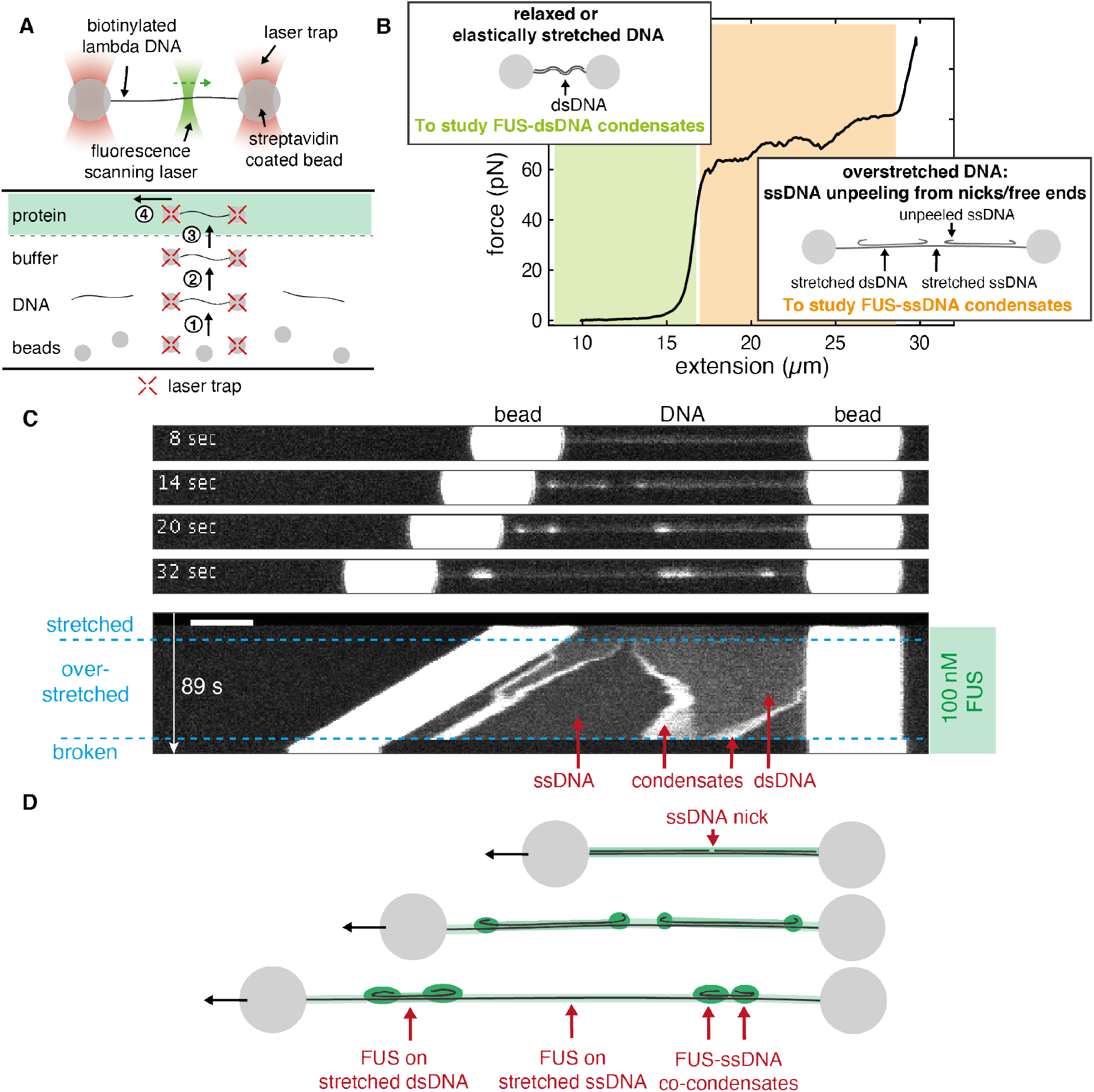
FUS forms co-condensates with ssDNA. (A) Schematics depicting the assembly and geometry of the optical tweezers-based assay. Lambda phage DNA is attached to Streptavidin-coated polystyrene beads held in optical traps. Binding of fluorescent FUS to DNA is recorded using scanning confocal fluorescence microscopy. Experiments are performed in a microfluidics flow chamber providing four separate experimental conditions via laminar flow fields. Steps for setting up the experiment in the flow chamber: (1) optical trapping of two Streptavidin coated polystyrene beads, (2) catching of a lambda phage DNA molecule that is biotinylated at its termini, (3) testing whether the tether is a single DNA molecule, (4) mechanical manipulation of the DNA in presence of FUS. (B) DNA mechanics and structure underlying our approach to study formation of FUS-ssDNA and FUS-dsDNA condensates. Investigation of FUS-dsDNA condensates is based on relaxed DNA. Investigation of FUS-ssDNA condensates is based on the gradual generation of unpeeled ssDNA during DNA overstretching. See also Movie S1 (C) Snap shots and kymograph showing FUS-DNA interaction and FUS-ssDNA condensate formation during overstretching of DNA at 100 nM FUS. Scale bar: 4 μm. (D) Schematics depicting DNA overstretching in presence of FUS. FUS homogeneously coats stretched ssDNA and dsDNA and forms condensates with unpeeled relaxed ssDNA.

To study how FUS interacts with single-stranded DNA (ssDNA) and double-stranded DNA (dsDNA), we exposed FUS to lambda phage DNA in different mechanical and structural states. The relationship between mechanical and structural properties of DNA is reflected in its force-extension curve (Figure 1B) (Smith, Cui and Bustamante, 1996; van Mameren *et al.*, 2009; Gross *et al.*, 2011). At extensions (*i.e.,* end-to-end distances) of up to about 0.9 times the contour length of the molecule (16.5 μm for lambda phage DNA) and at forces below ~10 pN, DNA behaves as an entropic spring. We refer to this regime as ‘relaxed’. At higher forces and at extensions that are similar to the contour length, the DNA molecule behaves like a Hookian spring. At extensions significantly higher than the contour length, the DNA molecule is ‘overstretched’. In the overstretching regime, a progressive increase of the end-to-end distance of the molecule results in a progressive conversion of dsDNA to ssDNA while DNA tension remains constant at around 65 pN. In this process, ssDNA is unpeeled, starting at free ssDNA ends. Free ends exist at nicks in the DNA backbone and at the ends of the dsDNA molecule. The overstretched DNA molecule consists of three distinct structural types of DNA: sections of stretched dsDNA interspersed with sections of stretched ssDNA (both load-bearing and at tensions of ~65 pN), with unpeeled and protruding ssDNA at the interfaces (Figure 1B, insets). The ratio between dsDNA and ssDNA is defined by the end-to-end distance to which the DNA molecule is overstretched. In this work, we used relaxed dsDNA to study the formation of FUS-dsDNA co-condensates, and we made use of unpeeled ssDNA protruding from overstretched DNA to study the formation of FUS-ssDNA co-condensates.

### FUS forms co-condensates with ssDNA

To first investigate the interactions of FUS with ssDNA, we used optical traps to hold in place a single lambda phage DNA molecule extended to its contour length of 16.5 μm and transferred into a microfluidics channel containing 100 nM FUS. Subsequently, we progressively increased its end-to-end distance to induce overstretching.

We observed that FUS attached to DNA in a spatially homogeneous manner upon transfer of the DNA molecule to the FUS channel (Figure 1C, Movie S1). When the DNA end-to-end distance was increased to achieve overstretching, the originally homogenous coverage of DNA by FUS became interspaced by regions that exhibited lower fluorescence intensity. At the interface between regions of higher and lower FUS intensity, FUS puncta emerged. When we increased the DNA end-to-end distance further, the length of regions with higher intensity decreased while the length of lower intensity regions increased. Concomitantly, the FUS puncta at the region interfaces grew in FUS intensity. Regions with high FUS intensity correspond to FUS unspecifically bound to stretched dsDNA (Figure 1D). Regions with low intensity correspond to FUS bound to stretched ssDNA, as these appear only during overstretching and grow with progressing overstretching (see Figure S2 for binding curves of FUS on stretched ssDNA and dsDNA). We interpret FUS puncta at interfaces between the low- and high density FUS regions as co-condensates of FUS with ssDNA, and provide evidence for condensation in the following sections. As the DNA is progressively overstretched, more and more unpeeled ssDNA is available, leading to growth of FUS-ssDNA co-condensates. We conclude that during overstretching, FUS binds to DNA in a manner that depends on the structural state of DNA: it homogeneously binds to dsDNA and ssDNA under tension, and forms condensates together with unpeeled ssDNA that is not under tension.

### FUS-ssDNA co-condensate formation is reversible

In what follows, we set out to study if FUS-ssDNA co-condensates recapitulate typical dynamic properties of biomolecular condensates observed *in vivo*. We first investigated the reversibility of the formation of FUS-ssDNA co-condensates. To test if FUS-ssDNA co-condensates can be dissolved by the removal of ssDNA, we performed a repetitive stretch-relax experiment consisting of two subsequent overstretch-relaxation cycles. The approach was based on the rationale that overstretching progressively generates free and unpeeled ssDNA available for co-condensation, while relaxation progressively removes it. We first overstretched a DNA molecule in presence of 100 nM FUS, by increasing its extension from 17 to 21 μm at a speed of 0.1 μm/s. The molecule was then relaxed again, followed by a second overstretch cycle. We recorded the spatiotemporal distribution of FUS along the entire molecule throughout the process (Figure 2A, Movie S2). In the example shown, we observed the formation of a condensate originating from a nick and a free terminal end on the right hand-side of the DNA molecule during the first overstretch. The size and brightness of condensates increased with progressive overstretching, in agreement with the findings presented in Figure 1B. During the subsequent relaxation cycle, the size and brightness of condensates decreased progressively until they completely disappeared. Notably, condensates formed at the precisely same locations and with essentially the same dynamics during the second overstretching cycle as they did during the first one. We conclude that FUS-ssDNA co-condensates can be dissolved by removal of available ssDNA.

**Figure 2.**
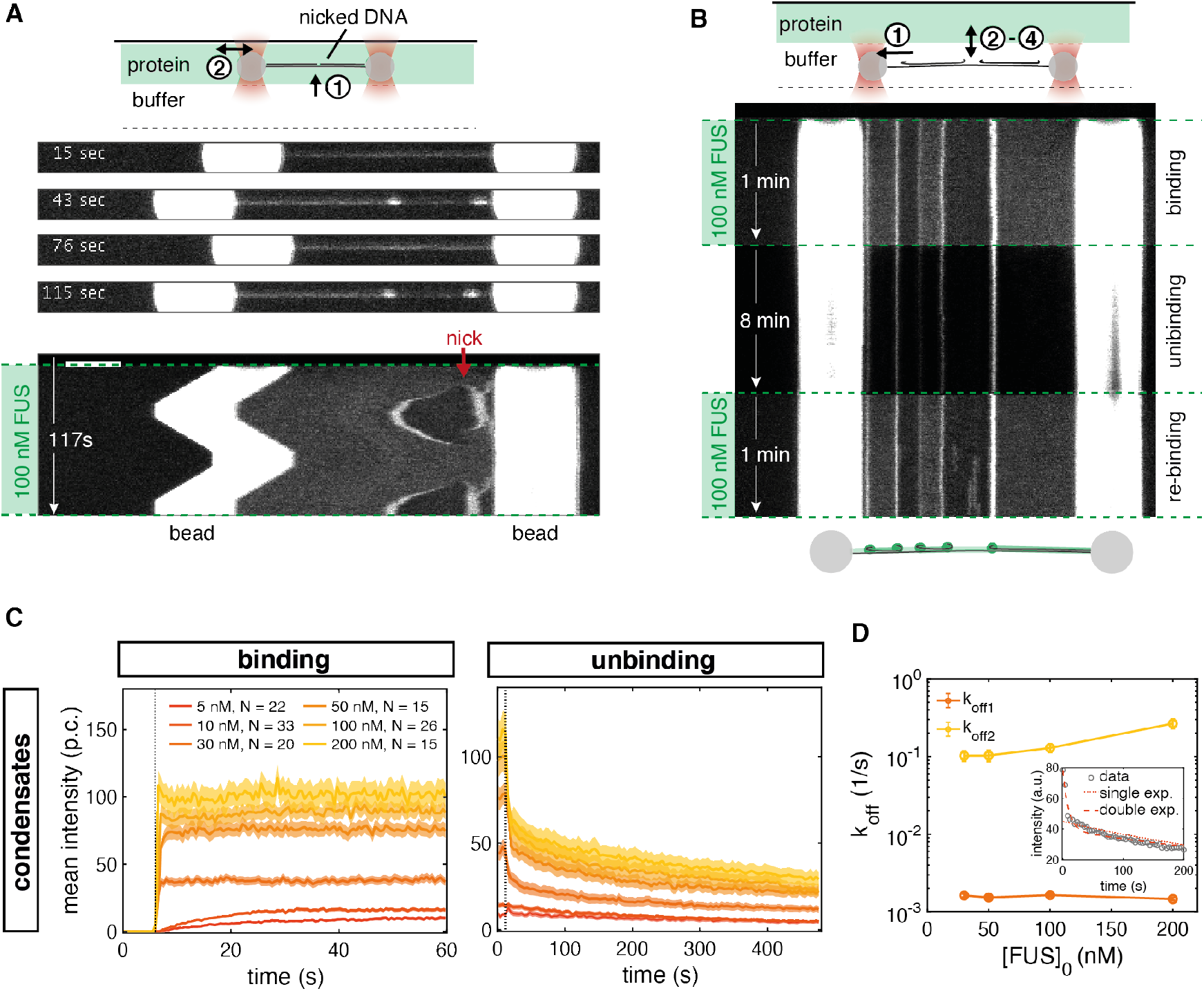
FUS-ssDNA co-condensate formation is reversible. (A) Snap shots and kymograph of repetitive overstretching experiments showing reversibility of condensate formation with respect to availability of a ssDNA scaffold. Scale bar: 4 μm. See also Movie S2 (B) Representative kymograph showing reversibility of FUS-ssDNA condensate formation with respect to availability of FUS tested in buffer exchange experiments. Condensates formed rapidly upon exposure of overstretched DNA to 100 nM FUS, slowly dissolved upon removal of free protein and rapidly re-formed upon re-exposure to free FUS. (C) Intensity-time traces of FUS-ssDNA condensates at different FUS concentrations during the binding and the unbinding step of the buffer exchange experiment; p.c. denotes the photon count. Plotted: mean ± STD. (D) Analysis of unbinding rates for condensates formed at different initial FUS concentrations. Inset: Intensity-time traces fitted with single and double exponentials. Fitting of intensity-time traces with double exponentials yielded 2 typical unbinding time scales in the range of seconds and hundreds of seconds, indicating at least 2 different interaction modes of FUS involved in FUS-ssDNA condensate formation. Error bars: 95 % confidence intervals.

To study if FUS-ssDNA co-condensates can be dissolved by the removal of free FUS from the environment, we performed binding-unbinding experiments by first overstretching a DNA molecule to 20 μm extension in absence of free FUS protein before moving it into (‘binding’), out of (‘unbinding’) and again into (‘re-binding’) the FUS protein channel (Figure 2B). We observed that in the binding process and upon entering the protein channel with 100 nM FUS, co-condensates rapidly formed, with a time scale that was below the temporal resolution of our imaging setup (0.5 s). Condensate formation was less rapid at lower concentrations of FUS (Figure 2C). In the unbinding process and in absence of free FUS protein, the size and brightness of condensates decreased progressively. However, within 480 s of observation time they did not disappear completely. Notably, the intensity-time traces of condensate dissolution deviated from simple single-exponential behavior, indicating that multiple types of interaction might be involved in stabilization of FUS-ssDNA co-condensates (Figure 2D). Upon re-exposure to free FUS protein during re-binding, condensates rapidly assumed the same size and intensity they had assumed in the initial binding step. Taken together, we conclude that FUS-ssDNA co-condensates dissolve when either ssDNA or free FUS is removed. FUS-ssDNA co-condensates form reversibly, which a) is indicative of a significant amount of protein turnover in these condensates, b) demonstrates that FUS-ssDNA interactions are key for co-condensation and c) demonstrates that FUS-FUS interactions, if they exist in these co-condensates, are not sufficient for maintaining a condensate in absence of ssDNA.

### FUS-ssDNA co-condensates are viscous droplets with liquid-like properties

Biomolecular condensates often show properties of liquid-like droplets *in vivo*. They deform under shear stress and can exhibit shape relaxation driven by surface tension (Brangwynne *et al.*, 2009; Jawerth *et al.*, 2018, 2020). We next investigated whether FUS-ssDNA co-condensates formed *in vitro* recapitulate this behavior. We first studied how these condensates react to the exertion of external mechanical perturbations. For that we increased the end-to-end distance of the DNA and hence the extend of overstretching in an abrupt and step-wise manner (steps every 10 s). This step-wise increase of the end-to-end distance within the overstretching regime instantaneously increases the amount ssDNA substrate available for co-condensate formation and causes the condensates to move with the propagating unpeeling front.

At 5 nM FUS, small FUS-ssDNA co-condensates emerged from the ends of the DNA molecule, which appear to instantaneously follow the propagation of unpeeling fronts (Figure 3A, left side). When increasing the amount of overstretch in a step-wise manner, condensates also grew in a step-wise fashion. This indicates that relaxation times are fast, and below the 1s interval between confocal image recordings. However, at 100 nM FUS, we observed that FUS-ssDNA co-condensates followed the step-wise bead movement with time delay and in a smooth, creeping-like manner, reminiscent of viscous droplet being dragged along a string (Figure 3A, right side, Movie S3). Leading and lagging edge of the condensates followed the bead movement on different response times, resulting in elongated condensate shapes. Elongated condensates relaxed towards more round shapes within the waiting time between steps (10s). This behavior is consistent with a viscoelastic response time of condensates associated with condensate viscosity and surface tension. We conclude that FUS-ssDNA co-condensates formed at concentrations of ~100 nM FUS display viscous material properties and exhibit viscoelastic shape relaxation.

**Figure 3.**
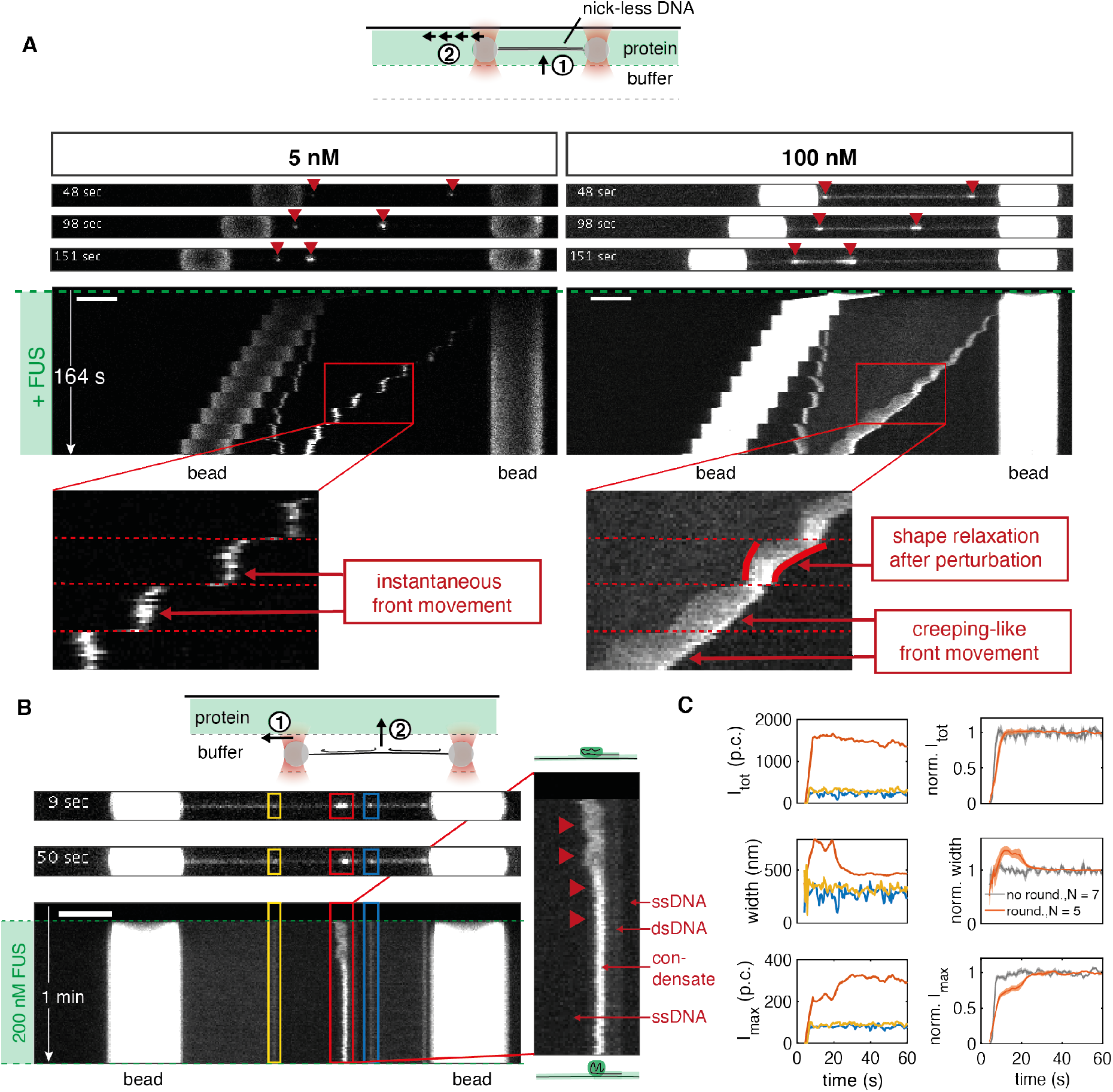
FUS-ssDNA co-condensates are viscous droplets with liquid-like properties. (A) Representative snap shots and kymographs of FUS on DNA molecules overstretched in a step-wise manner in presence of different FUS concentrations. At 5 nM FUS, condensates had a point-like morphology and instantaneously grew and moved along the DNA when the DNA end-to-end distance was increased (left side, zoom). At 100 nM FUS, condensates grew and moved along the DNA in a creeping-like manner when the DNA end-to-end distance was increased. They elongated and showed shape relaxations on slow time scales compared to the fast-imposed external perturbations, reminiscent of viscous, liquid-like droplets (right side, zoom). (B) Representative snap shots and kymographs of a binding experiment performed at 200 nM FUS. Occasional shape changes from an initial elongated to a rounded morphology were observed (red condensate and zoom). (C) Left side: quantification of shape changes of the example condensates. From top to bottom: total intensity, width and maximum intensity of individual condensates over time. While the total intensity of the red condensate remained constant over the course of the experiment, its width decreased while its maximum intensity increased until they levelled off. Right side: quantification of shape changes of condensate ensemble. From top to bottom: normalized total intensity, normalized width and normalized maximum intensity of condensates over time. At 200 nM FUS, 5 out of 12 condensates showed rounding, decreasing their width to 70 % of their initial with and increasing their maximum intensity to 140 % of their initial intensity while keeping their total intensity constant within about 20 s. Traces show mean ± SEM, p.c. denotes the photon count.

We next set out to find additional signatures for viscoelastic shape relaxation of FUS-ssDNA co-condensates. To this end we investigated condensate shape changes after their formation. Figure 3B presents snapshots and the kymograph of a typical binding experiment performed at 200 nM FUS, showing how FUS assembles on the different segments of the overstretched DNA molecule upon exposure to FUS. In the representative example shown, while the two small condensates (marked in blue and yellow) did not change their shape after formation, the big condensate (marked in red) transitioned from an initially elongated towards a round shape within ~20 s (Figure 3C). We conclude that FUS-ssDNA co-condensates display viscoelastic shape relaxations on a timescale that is of the order of 10 s. We have thus revealed two types of shape relaxation of FUS-ssDNA co-condensates consistent with liquid-like behavior: they deform upon external mechanical perturbations and they relax their shape after rapid formation.

### FUS associating with ssDNA generates a sticky FUS-ssDNA polymer

We speculate that FUS can form dynamic co-condensates with ssDNA because the association of FUS with ssDNA generates a self-interacting polymer which undergoes a globular collapse to form a liquid-like FUS-DNA co-condensate (Halperin and Goldbart, 2000; Polotsky *et al.*, 2010; Cristofalo *et al.*, 2020). Here, FUS-FUS or additional FUS-DNA interactions could act like a ‘molecular glue’ when two FUS-coated ssDNA fragments meet, which would prevent their dissociation. To test if FUS-DNA indeed behaves like a sticky polymer, we overstretched single DNA molecules whose top strands were by chance nicked at certain locations. We refer to the single strand of the dsDNA molecule that remains physically attached to the two polystyrene beads as the “principal strand”, while the complementary strand which becomes progressively unpeeled during overstretching is referred to as the “top strand”. When a dsDNA molecule with a nicked top strand is overstretched to completeness (to 1.7 times its contour length), the unpeeled top strand fragments should dissociate and detach completely from the principal strand. We here tested if the interaction between FUS and ssDNA could interfere with this top strand detachment process.

Figure 4A (left side) shows the kymograph of a typical stepwise overstretching experiment performed at 5 nM FUS. We observe ssDNA unpeeling and condensation of ssDNA fragments with FUS, originating from the two terminal ends of the DNA molecule and from two nicks. When two unpeeling fronts met, they fused and subsequently disappeared from the field of view. This indicates that the corresponding ssDNA top strand fragment completely detached from the principal strand. Notably, all three ssDNA top strand fragments dissociated from the principal strand, but the principal strand was still intact after dissociation of the last top strand fragments. However, in the example kymograph for the experiment performed at 100 nM FUS (Figure 4A, right side), the top strand fragments did not fall off after unpeeling fronts of the individual fragments met in the course of overstretching. Rather, the top strand fragments remained attached to the principal strand. Taken together, our observations are consistent with the picture that FUS-coated ssDNA behaves like a sticky polymer, which serves to hold isolated fragments of ssDNA attached to regions of dissociation.

**Figure 4.**
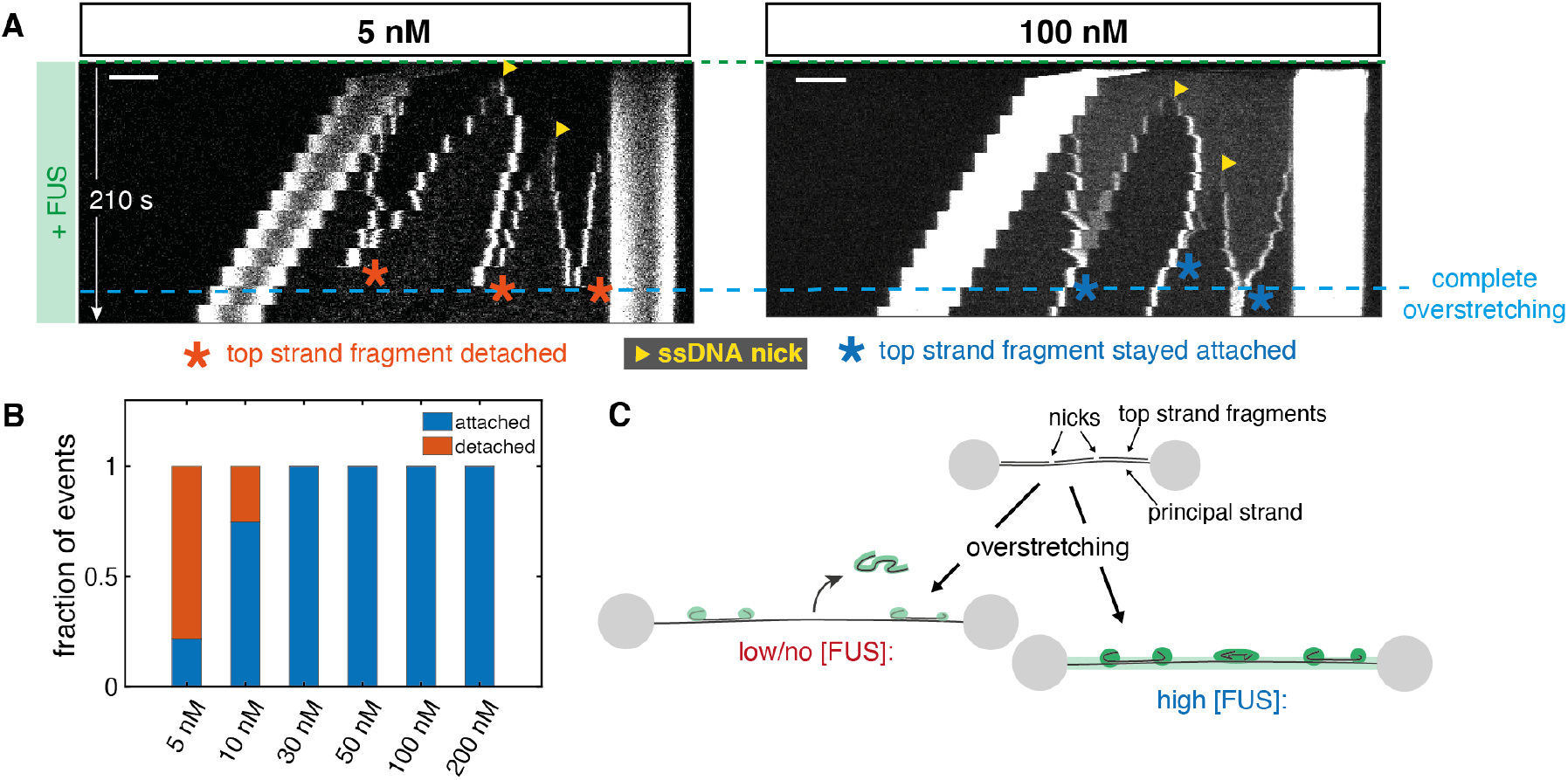
FUS associated with ssDNA generates a sticky FUS-ssDNA polymer. (A) Representative kymographs showing the influence of FUS-ssDNA interaction on the dissociation of DNA fragments when the DNA molecules are overstretched. The principal strand is the single strand of the dsDNA molecule attached to the beads. At 5 nM FUS, fragments generated by overstretching detached from the principal strand while at 100 nM FUS, fragments stayed attached to the principal strand. Scale bar: 4 μm (B) Quantification of the fraction of fragments that detached from the principal strand vs. the fraction of fragments that stayed attached in step-wise overstretching experiments. Only at 5 and 10 nM FUS, fragments were able to detach from the principal strand, while FUS-ssDNA condensates formed at higher FUS concentrations always stayed attached. Number of events: 5 nM: 23, 10 nM: 16, 30 nM: 14, 50 nM: 7, 100 nM: 23, 200 nM: 18. (C) Illustration of the fragment detachment/attachment process.

If self-interactions of the FUS-ssDNA polymer arise from FUS-FUS interactions, or from FUS-ssDNA interactions that are in addition to the normal mode of association of FUS to ssDNA, we would expect that these self-interactions depend on FUS concentration. We analyzed unpeeling events from experiments performed in the concentration range between 5 and 200 nM FUS, and classified them into “detached” (a top strand fragment disappeared from the principal strand when two corresponding unpeeling fronts met while the principal strand stayed intact) and “attached” (a top strand fragment remained attached to the principal strand when two corresponding unpeeling fronts met). We found that only for FUS concentrations below 30 nM, a considerable fraction of unpeeled top strand fragments detached from the principal strand (Figure 4B), while they remained associated at higher concentrations. We conclude that self-interactions of the FUS-ssDNA polymer depend on FUS concentration. FUS-ssDNA co-condensate have liquid-like properties (see above). Capillary forces are mechanical forces that are generated by a fluid when contacting a surface (de Gennes, Brochard-Wyart and Quéré, 2004; Quail *et al.*, 2020). Given that self-interactions of the FUS-ssDNA polymer can generate a liquid phase, it is tempting to speculate that this can give rise to generalized capillary forces for liquid phases consisting of collapsed self-interacting polymers. For FUS, these could arise when the liquid phase contacts other FUS-coated DNA strands, resulting in the continued adhesion of condensates with principal strands in the experiments described above, and delaying force induced disruption of dsDNA strands (Figure S3). It is tempting to speculate that these behaviors are related the ability of FUS-dsDNA interactions to act as a molecular glue in the context of the DNA damage response. This is interesting as one might expect that an immediate response to DNA damage requires prevention of DNA fragments from leaving the damage site.

So far, we have shown that FUS forms dynamic co-condensates with ssDNA and that these condensates show various properties that are also typical for protein-nucleic acid-based organelles observed *in vivo*: their formation is reversible, they exchange constituents with the environment and they show liquid-like material properties. Co-condensation also mediates stickiness and the adhesion of separate ssDNA strands. We next used the possibilities offered by our single molecule manipulation approach to reveal the physicochemical mechanisms underlying the formation of such FUS-ssDNA condensates.

### FUS-ssDNA co-condensation is based on FUS adsorbing in a single layer on ssDNA

We were interested to understand if ssDNA in FUS-ssDNA condensates is coated with a single adsorption layers of FUS with every FUS molecule directly binding to ssDNA, or if multiple layers of FUS are present with some FUS molecules not directly bound to ssDNA. We first investigated how the size of FUS-ssDNA co-condensates depends on the number of incorporated nucleotides. For this we utilized the step-wise overstretching assay introduced in Figures 3 and 4. By controlling the end-to-end distance of the DNA molecule within the overstretching regime in a step-wise manner, we controlled the total number of unpeeled ssDNA nucleotides available for FUS-ssDNA co-condensate formation (Figure 5A). By utilizing nick-free DNA molecules only, we ensured that ssDNA unpeeling during overstretching only occurred from the two ends of the DNA molecules. By measuring the distance between each of the two forming condensates and the respective beads, and taking into account the length of a single nucleotide under the applied tension of around 65 pN (0.58 nm, (Smith, Cui and Bustamante, 1996; van Mameren *et al.*, 2009; Gross *et al.*, 2011)), we were able to determine the number of nucleotides available for incorporation into each of the two FUS-ssDNA co-condensates (see Experimental Procedures for details). Further, we determined the integrated FUS fluorescence intensity associated with each condensate. Notably, we calibrated the FUS fluorescence intensity to arrive at a number of FUS-EGFP molecules in the condensate, using a calibration procedure that relied on individual dCas9-EGFP molecules tightly bound to lambda phage DNA molecules (see Experimental Methods and Figure S4) (Morin *et al.*, 2020). We found that at all FUS concentrations investigated (between 1 nM and 200 nM FUS), the number of FUS molecules in a condensate was proportional to the number of incorporated nucleotides, with a slope that depends on the FUS concentration (Figure 5B). This confirms that a) the number of FUS molecules in a FUS-ssDNA co-condensate is determined by the amount of available ssDNA substrate, and that b) co-condensate stoichiometry (*i.e*., the ratio between number of proteins and number of nucleotides in a condensate) is independent of the size of the condensate, as is expected for co-condensation. More precisely, co-condensate stoichiometry is independent of the total number of ssDNA nucleotides in the co-condensate but depends on bulk FUS concentration (Figure 5B). The ratio between the number of proteins and the number of nucleotides (nt) in a co-condensate (*i.e.,* the slopes of the relations in Figure 5B) informs about the degree of ssDNA substrate occupation by FUS. This ratio increased with increasing FUS concentrations between 1 and 50 nM, and saturated at higher concentrations (Figure 5C). Strikingly, this saturation curve was well described by a simple Langmuir adsorption model (*K_d_* = 31.5 nM (11.3 - 51.8 nM) (numbers in brackets indicate the lower and upper bound of the 95% confidence interval, unless otherwise noted), saturation level p_0_ = 0.08 molecules/nt (0.06 - 0.10 molecules/nt) for FUS-recruitment to ssDNA in FUS-ssDNA co-condensates). This model assumes that ligands occupy binding sites on the substrate independently and with negligible ligand-ligand interactions (Langmuir, 1918; Mitchison, 2020). Furthermore, the saturation level of the Langmuir adsorption curve (Figure 5C) implies a saturated density of FUS on ssDNA with approximately one FUS molecule every 12.4 nucleotides of ssDNA. Taken together, our data suggests that FUS in FUS-ssDNA co-condensates forms a single adsorption layer on ssDNA, with every FUS molecule directly bound to ssDNA. Binding occurs without detectable cooperativity despite the fact that FUS-FUS interactions within such a FUS-ssDNA co-condensate appear to collectively generate the capillary forces that drive co-condensation and condensate shape changes.

**Figure 5.**
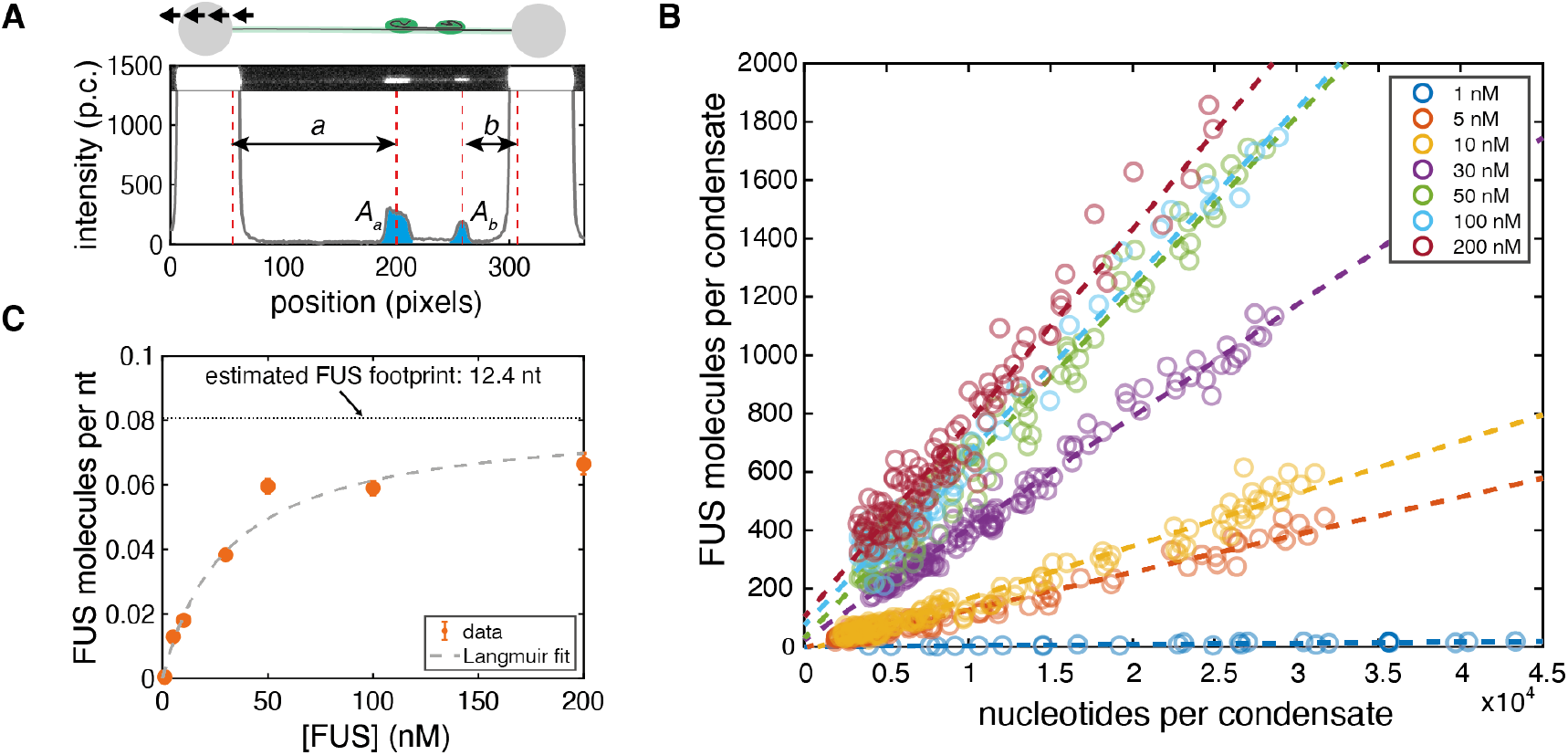
FUS-ssDNA co-condensation is based on FUS adsorbing in a single layer on ssDNA. (A) Intensity of FUS-ssDNA condensates and the number of potentially incorporated nucleotides were extracted from step-wise overstretching experiments (p.c. denotes photon count). *A_a_* and *A_b_* are integrated intensities of condensates, *a* and *b* are the pieces of ssDNA incorporated in each of them. (B) Number of FUS molecules vs. number of nucleotides incorporated in each condensate. Number of events: 1 nM: 25, 5 nM: 72, 10 nM: 68, 30 nM: 69, 50 nM: 47, 100 nM: 38, 200 nM: 59. An event is a single condensate observed during a single stretching step in a step-wise overstretching experiment. Dashed lines: linear fits to data points at the corresponding FUS concentration. Intensities were converted into numbers of FUS molecules by calibration with single dCas9-GFP molecules (Figure S4). (C) Number of FUS molecules per nucleotide in condensates vs. FUS concentration obtained from linear fitting in (B). Data is fitted by a Langmuir binding isotherm, implying that the Langmuir-like recruitment of a monolayer of FUS to ssDNA underlies FUS-ssDNA condensate formation. The saturation value of the curve (dotted horizontal line) indicates a footprint of the FUS molecule inside FUS-ssDNA condensates of 12.4 nucleotides. Plotted: orange: result of linear fitting in (B) within 95 % confidence intervals. Grey dashed lines: Langmuir fit.

### FUS monolayer adsorption to dsDNA and LCD mediated interactions lead to FUS-dsDNA co-condensate formation

Given that FUS does not only have an affinity for single- but also for double-stranded DNA (Figure 1C), we next investigated whether FUS can also form co-condensates with dsDNA. For this, we attached a single dsDNA molecule to a Streptavidin-coated bead held in an optical trap and applied an external buffer flow to stretch the DNA. We then moved the stretched bead-DNA construct to a channel containing 100 nM FUS while the flow was maintained (Figure 6A). When moving the flow-stretched dsDNA molecule into the protein channel, we observed that a) the dsDNA molecule became immediately coated with FUS (Figure 6B, Movie S4) and b) a co-condensate appeared to form at the free end of the dsDNA molecule, rapidly moving towards the bead and increasing in size with decreasing distance to the bead. Co-condensation was abolished when the low-complexity domain of FUS was not present (Figure 6C), indicating that, as expected, the low-complexity domain plays a role in mediating the FUS-FUS interactions necessary for co-condensation of FUS with dsDNA. Together, this provides evidence that the interaction of FUS with dsDNA leads to the formation of a FUS-dsDNA co-condensate even in presence of DNA tension.

**Figure 6.**
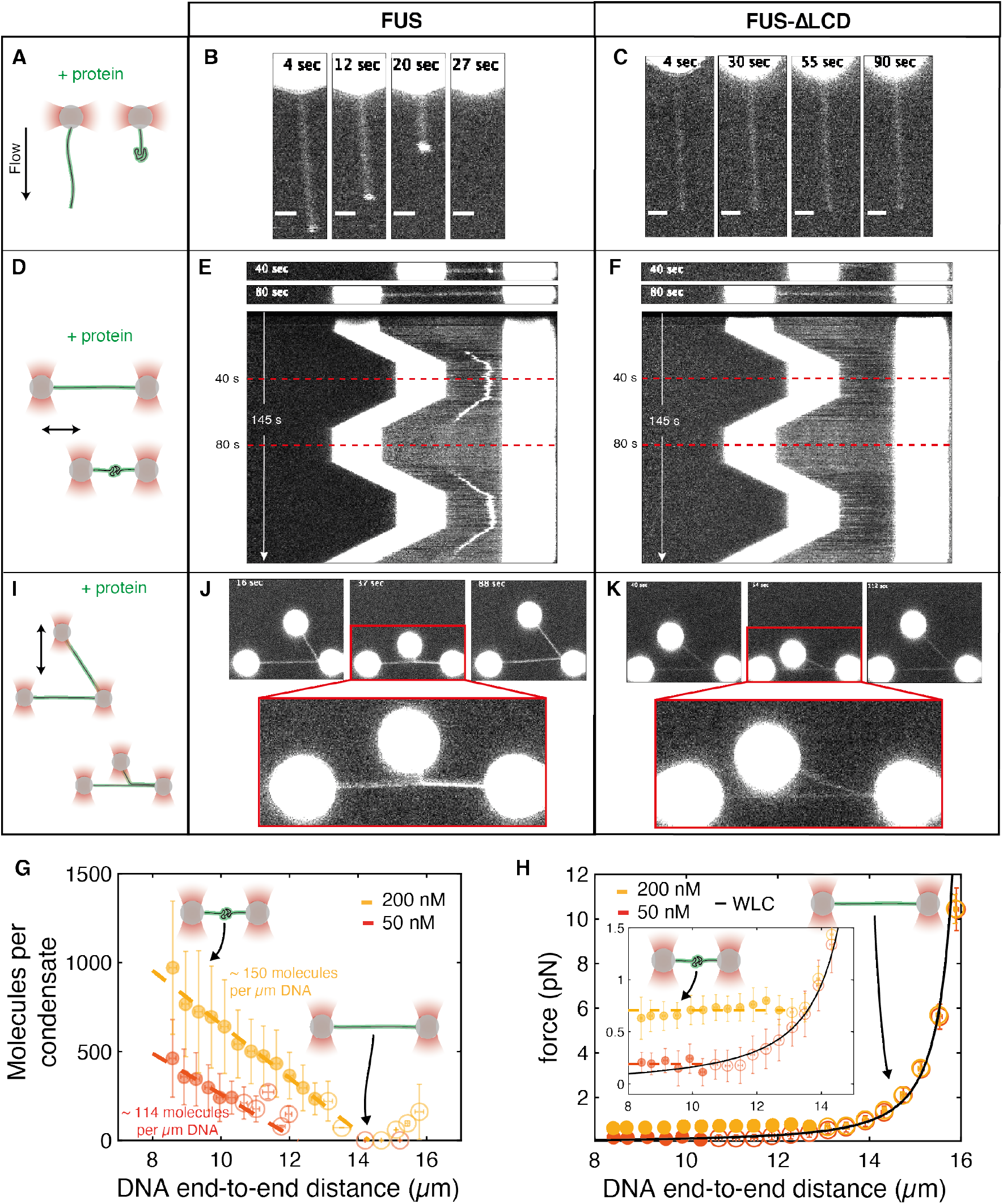
FUS monolayer adsorption on dsDNA and low-complexity domain mediated interactions lead to FUS-dsDNA co-condensate formation. Representative snap shots and kymographs of FUS-dsDNA interaction assessed in 3 optical tweezers-based assays. (A) Individual dsDNA molecules were attached via one end to trapped beads and stretched by flow in presence of protein. (B) FUS homogeneously adsorbs on and condenses hydrodynamically stretched dsDNA. (C) FUS with deleted low-complexity domain (FUSΔLCD) homogeneously binds to hydrodynamically stretched dsDNA, but does not mediate condensation. (D) dsDNA attached to two trapped beads via both ends was stretched and relaxed between 8 and 16 μm end-to-end distance in presence of protein. (E) FUS forms reversible condensates with relaxed dsDNA. (F) FUSΔLCD homogeneously binds to dsDNA, but does not mediate condensation. (G) Number of FUS molecules in FUS-dsDNA condensates vs. DNA end-to-end distance studied in dual-trap optical tweezers experiments. Data was obtained from tracking the condensate intensity and DNA end-to-end distance during the initial relaxation from 16 to 8 μm DNA extension. Number of FUS molecules inside condensates was estimated from condensate intensity using the calibration procedure described in Figure S4. Number of FUS molecules inside condensates linearly increases with decreasing DNA end-to-end distance, while the slope of this increase depends on the FUS concentration and hence on the FUS coverage of dsDNA. Condensate formation only occurs below a critical, FUS concentration dependent DNA end-to-end distance *L_crit_* (see Figure S5D). Red: condensates formed at 50 nM FUS (29 individual DNA molecules); yellow: condensates formed at 200 nM FUS (22 individual DNA molecules). Filled circles: data points classified as ‘condensate’; open circles: data points classified as ‘no condensate’ (classification with respect to *L_crit_*). Mean ± STD. Dashed lines: linear fits indicating the linear increase of condensate size with decreasing DNA end-to-end distance. (H) DNA tension vs. DNA end-to-end distance measured in dual trap experiments. When DNA is relaxed in presence of FUS starting from 16 μm end-to-end distance, the measured relationship between force and DNA end-to-end distance coincides with the one of ‘naked’ DNA (black line, Worm-like Chain model (WLC)). Below the critical DNA end-to-end distance, the force remains constant when the end-to-end distance is reduced further. Red: condensates formed at 50 nM FUS (29 individual DNA molecules); yellow: condensates formed at 200 nM FUS (22 individual DNA molecules). Filled circles: data points classified as ‘condensate’; open circles: data points classified as ‘no condensate’ (classification with respect to *L_crit_*). Mean ± STD. Dashed lines: linear fits indicating the force buffering by condensates when DNA end-to-end distance is decreased. (I) Two dsDNA molecules were attached to three trapped beads in an L-like configuration. One bead was moved to approach the molecules and hence to allow for protein mediated DNA zippering. (J) FUS mediates capillary-like forces between dsDNA strands. (K) DNA zippering is lost by deletion of the FUS LCD. Scale bars: (B), (C): 2 μm; (E), (F), (J), (K): bead diameter 4 μm

**Figure 7.**
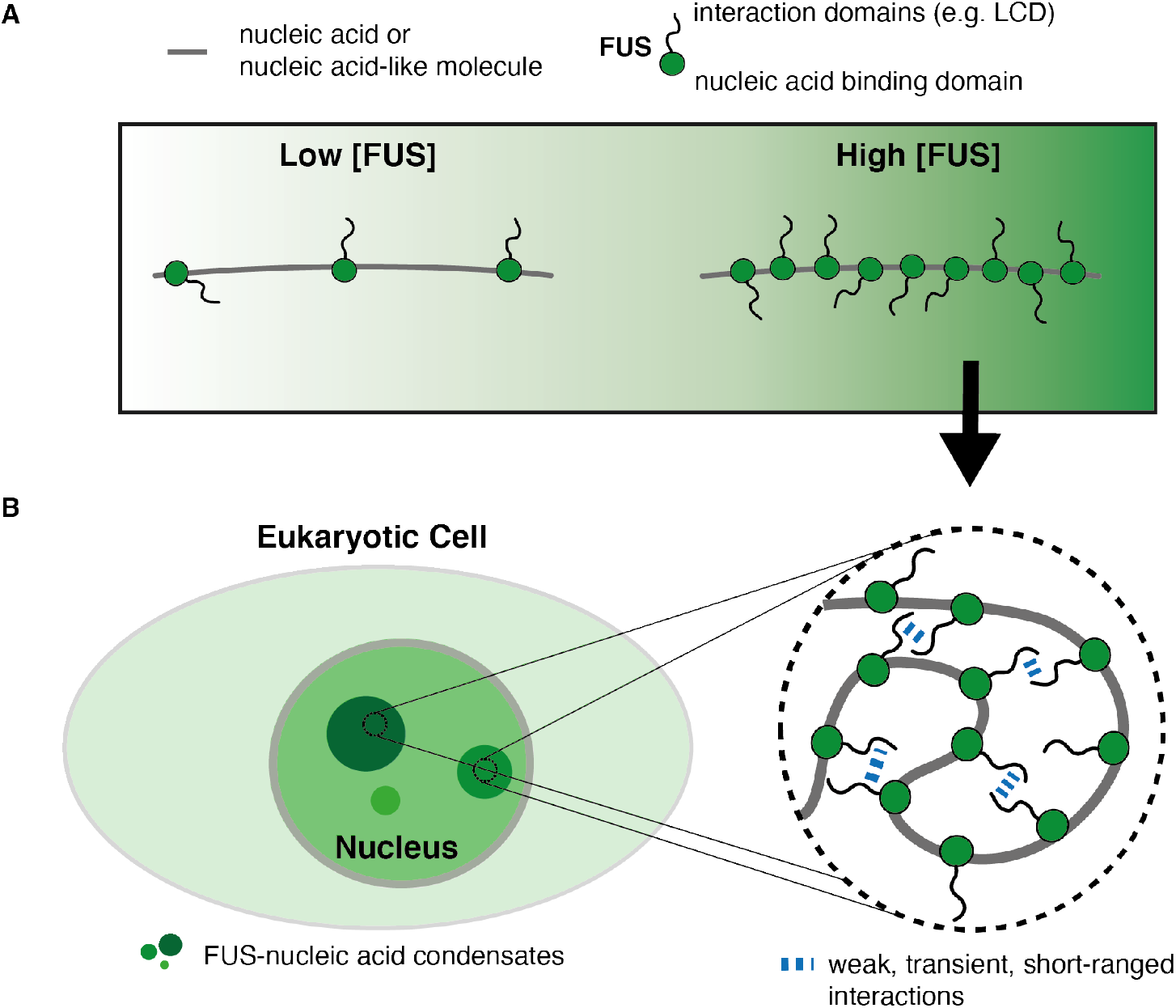
Biomolecular condensate formation based on monolayer protein recruitment to nucleic acids. (A) Nucleic acids or nucleic acid-like polymers recruit monolayers of proteins. (B) Protein adsorption on nucleic acids gives rise to an effective self-interacting protein-nucleic acid polymer. Collapse of this self-interacting polymer leads to the formation of protein-nucleic acid co-condensates reminiscent of biomolecular condensates observed in cell nuclei.

To better investigate the co-condensation process, we next attached a dsDNA molecule to two beads held in place in optical traps, and repeatedly relaxed and stretched the molecule between 8 μm and 16 μm end-to-end distance and thus to a length slightly below its contour length in a solution containing 200 nM FUS (Figure 6D). Again, we observed that FUS assembled homogeneously on the stretched dsDNA molecule (Figure 6E, Movie S5). Strikingly, a single FUS-DNA co-condensate emerged when the DNA was relaxed to an end-to-end distance below ~14 μm, which grew in FUS amount with decreasing DNA end-to-end distance. The condensate dissolved again when the DNA was stretched beyond ~14 μm, and it re-formed with similar dynamics when the DNA was relaxed again, albeit at a slightly different position. Again, condensate formation depended on the presence of the low-complexity domain of FUS (Figure 6F). To conclude, FUS can form dynamic, reversible co-condensates with relaxed dsDNA.

We next asked whether these FUS-dsDNA co-condensates indeed form a separate physical phase. We draw an analogy to the phase transition between liquid water and vapor (Atkins, de Paula and Keeler, 2017) when a pot of water is put onto a hot stove, the temperature of the water will not surpass 100°C. Instead of increasing the temperature, energy input will cause water to transition from the liquid phase to the vapor phase while the temperature remains constant. This analogy is helpful for understanding the dissolution of FUS-dsDNA co-condensates by mechanically extracting FUS coated dsDNA from the condensate. We predict two effects to occur when the end-to-end distance of a FUS coated, condensed dsDNA molecule is increased. First, mass conservation implies that as FUS coated DNA is progressively extracted from the condensate, the amount of material in the FUS-dsDNA co-condensate should decrease by corresponding amounts. Second, the dissolution of FUS-dsDNA co-condensates should occur at a constant DNA tension, similar to the constant temperature observed for the transition between liquid and gaseous water (Cristofalo *et al.*, 2020; Quail *et al.*, 2020).

To test the first prediction, we used the dual-trap experiment to form and dissolve FUS-dsDNA co-condensates at 50 and 200 nM FUS. Mass conservation implies that the number of FUS molecules inside a condensate increases proportionally with the amount of co-condensing dsDNA. In other words, the amount of FUS in the co-condensate should increase linearly with decreasing DNA end-to-end distance, which is what we observed (Figure 6G). Furthermore, the absolute value of the slope of the linear relationship between number of FUS molecules in a co-condensate and the DNA end-to-end distance increased with increasing FUS concentration (Figure 6G). At 50 nM FUS, ~114 FUS molecules are bound per μm of DNA in a co-condensate (corresponding to a spacing of one FUS molecule every ~26 bp), while at 200 nM FUS ~150 FUS molecules are bound per μm of DNA in a co-condensate (corresponding to a spacing of one FUS molecule every ~20 bp) (also see Figure S2). This reveals that FUS adsorbs in a single layer on DNA at both concentrations investigated, with enough space between FUS molecules to allow each FUS molecule to directly bind to dsDNA. An analysis of the probability for co-condensate formation as a function of DNA end-to-end distance revealed a sharp transition at 10.5 μm (10.4 μm - 10.6 μm) at 50 nM FUS and 12.9 μm (12.7 μm - 13.1 μm) at 200 nM FUS (Figure S5D). This indicates that co-condensation occurs below a critical DNA end-to-end distance *L_crit_* that depends on FUS concentration. Taken together, we conclude that as FUS coated DNA is progressively extracted from the FUS-dsDNA co-condensate, the amount of material in the FUS-dsDNA co-condensate decreases by corresponding amounts.

We next tested the second prediction and investigated the range of DNA tensions at which FUS-dsDNA co-condensates form (Figure 6H and Figure S5G). Using the dual trap tweezer assay we found that as FUS-coated dsDNA is relaxed starting from an initially stretched configuration (16 μm end-to-end distance), the relation between force and DNA end-to-end distance follows the expected Worm-like Chain (WLC) behavior as long as its end-to-end distance is above *L_crit_*. Strikingly, when the end-to-end distance was reduced below *L_crit_* (and hence when a condensate forms), trap force remained constant (0.19 ± 0.05 pN at 50 nM FUS, 0.71 ± 0.05 pN at 200 nM FUS (mean ± STD)). Furthermore, condensates of various sizes coexisted at essentially the same DNA tension (Figure S5G). Note also that in the region where the WLC transitions into the constant force regime a slight dip in force was observed, indicative of a small but finite surface tension of the condensate. A theoretical description of protein-DNA co-condensation in the optical trap suggests that this dip corresponds to a surface tension of the order of 0.15 pN/μm (Figure S5F and supplementary experimental procedures). Together, this provides evidence that a first-order phase transition underlies the formation of FUS-dsDNA co-condensates.

We next set out to estimate the condensation free energy per FUS molecule (Quail *et al.*, 2020). At DNA end-to-end distances far below the critical DNA length and in the case of low surface tension, the constant force generated by the co-condensate reeling in DNA is determined by the condensation free energy per volume μ and the DNA packing factor ⍺. The packing factor is a measure for the scaling between length of condensed DNA and the volume of the condensate. We estimated ⍺ using the FUS concentration dependent FUS coverage of dsDNA inside condensates (slope in Figures 6G and S5E) and the molecular volume of FUS inside condensates *V_m_* (Figure S5F). We found that values of ⍺ (~ 0.05 μm^2^ at 50 nM FUS, ~ 0.06 μm^2^ at 200 nM FUS) were similar in magnitude to those reported for a DNA-protein phase containing the transcription factor FoxA1 (Quail *et al.*, 2020). The condensation free energy per volume obtained using the packing factors and corresponding critical forces was ~ 4.1 pN/μm^2^ at 50 nM FUS and ~ 11.9 pN/μm^2^ at 200 nM FUS. With a FUS density inside condensates of about 2500 molecules/μm^3^ (specified by the molecular Volume *V_m_*), this provides an estimate of the condensation free energies of ~ 0.4 kT/FUS at 50 nM FUS and ~ 1.1 kT/FUS at 200 nM FUS. Taken together, FUS adsorbing in a single layer on DNA effectively generates a sticky FUS-DNA polymer that can collapse to form a liquid-like FUS-DNA co-condensate. For double-stranded DNA, this condensation occurs at constant DNA tension which is a clear signature of a mesoscopic first-order phase transition.

We next set out to test if single layer adsorbed FUS can mediate adhesion of separate dsDNA strands. For that we attached two FUS-coated dsDNA strands to three beads in an L-like configuration, with DNA strands held at tensions above the critical tension for FUS-dsDNA condensate formation (Figure 6I). By moving one of the beads relative to the others, we were able to bring the two FUS-coated dsDNA molecules in close proximity in order to test if they adhere to each other. We found that the two FUS-coated dsDNA strands adhered to each other and “zippered up” at 100 nM FUS (Figure 6J, Movie S6). Zippering was reversed by pulling the DNA strands away from each other and re-established by moving DNA strands closer. Furthermore, zippering depends on the presence of the low-complexity domain (Figure 6H). Taken together, our data indicates that a single layer of FUS attached to DNA can mediate dynamic adhesion of separate DNA strands, opening up the possibility for this mechanism to be involved in long-range genome organization.

## Discussion

The discovery that membrane-less compartments can be formed by liquid-like biomolecular condensates and that phase separation can contribute to the spatiotemporal organization of intracellular biochemistry has opened up new perspectives in cell biology (Hyman, Weber and Jülicher, 2014; Banani *et al.*, 2017). Here, we have demonstrated that FUS molecules can adsorb in a single layer on DNA, which is well described by a Langmuir isotherm (Figures 5 and S2). At low dsDNA tension, self-interactions between FUS molecules facilitate co-condensation and formation of FUS-DNA co-condensates (Figure 6). Here, changing dsDNA extension shifts the balance between the co-condensate and the FUS-coated dsDNA molecule, and results in the co-condensate growing at the expense of stretched dsDNA. The process of co-condensation is a chemo-mechanical process that converts chemical potential changes to mechanical forces. These generalized capillary forces can exert tension on the DNA that remains outside the condensate. Growth of co-condensates occurs at constant DNA tension, consistent with a mesoscopic first order phase transition as is expected for a physical condensation process. We find that the constant tension depends on the FUS concentration and is of the order of 1 pN. For comparison, forces required for unfolding individual proteins typically are higher and in the range of tens of pN (Gupta *et al.*, 2016; Ganim and Rief, 2017; sen Mojumdar *et al.*, 2017). Also, the stall force of RNA Pol II is at least an order of magnitude higher, opening up the possibility that transcription can proceed essentially unhindered in the presence of such capillary forces (Yin *et al.*, 1995). Protein-DNA co-condensation involves the collective binding of many proteins to a DNA substrate. Here we demonstrate that upon single-layer binding the FUS coated DNA molecule undergoes co-condensation. In other scenarios, interactions of protein ligands and DNA surfaces could lead to multilayer-adsorption (Mitchison, 2020), or the formation of protein microphases via prewetting transitions (Morin *et al.*, 2020) (Figure S1).

We speculate that the mechanism we describe here is relevant for other processes of DNA compaction, such as heterochromatin formation driven by HP1⍺ (Larson *et al.*, 2017; Strom *et al.*, 2017; Larson and Narlikar, 2018; Sanulli *et al.*, 2019; Keenen *et al.*, 2021). We further suggest that generalized capillary forces arising in liquid-like co-condensates play an important role in other biological processes such as the transcription-dependent organization of chromatin (Cho *et al.*, 2018; Sabari *et al.*, 2018; Thompson *et al.*, 2018; Henninger *et al.*, 2021) (Figure S6A), the formation of viral replication compartments (Schmid *et al.*, 2014; Heinrich *et al.*, 2018; McSwiggen *et al.*, 2019; Nevers *et al.*, 2020) (Figure S6C), and the DNA damage response (Altmeyer *et al.*, 2015; Patel *et al.*, 2015; Aleksandrov *et al.*, 2018; Naumann *et al.*, 2018; Levone *et al.*, 2021) (Figure S6B). With respect to the latter, we have shown that pairs of FUS-coated DNA can bind together and exert adhesion forces onto each other (Figures 4 and 6) It is possible that inside the cell, such adhesion forces prevent DNA fragments to leave damage sites during DNA repair. An interesting question for future research is to understand if poly(ADP)ribose (PAR) triggers FUS-DNA co-condensation at the damage site, thereby preventing the escape of DNA damage fragments. Taken together, we suggest protein-nucleic acid co-condensation constitutes a general mechanism for forming intracellular compartments.

## Acknowledgments

We thank members of the S.W.G lab for fruitful discussion. S.W.G. was supported by the DFG (SPP 1782, GSC 97, GR 3271/2, GR 3271/3, GR 3271/4) and the European Research Council (grant 742712). R.R. and A.H. acknowledge support by the NOMIS foundation. A.H. is supported by the Hermann und Lilly-Schilling Stiftung für medizinische Forschung im Stifterverband. We thank Stefan Golfier for contributing the sgRNA and dCas9-EGFP for single molecule intensity quantification experiments. We further acknowledge K.M. Crell, S. Kaufmann and F. Thonwart for providing reagents. We also thank J. Brugues, A. Gladfelter, A. Hyman, T. Mitchison, W. Snead and T. Quail for discussions and critical comments on the manuscript.

---

**Figure S1.**
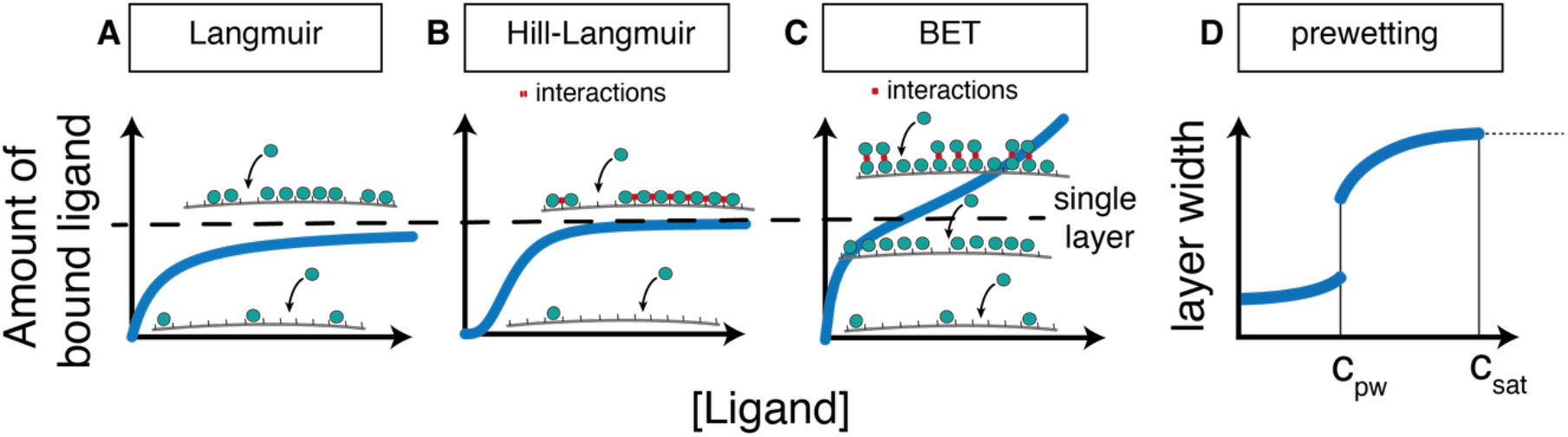
Ligand adsorption on substrates. The type of adsorption of ligands on a substrate depends in the relative strengths of ligand-ligand and ligand-substrate interactions. (A) If ligand-ligand interactions are negligible, the ligands form a single layer on the substrate, with a lattice site occupancy that increases with increasing ligand concentration and approaches saturation at high ligand concentrations (Langmuir model). (B) In presence of cooperative ligand-ligand interactions that support association with the substrate, the ligand occupancy of the scaffold follows a switch-like, sigmoidal trend. Increase of ligand concentration results in the formation of a single ligand layer on the scaffold (Hill-Langmuir model). (C) In presence of attractive ligand-ligand interactions, association of ligands to a substrate can be described using the BET model. Increase in ligand concentration first leads to the formation of a single protein layer on the substrate and later to the formation of multiple layers of ligands on top of the initial layer. In contrast to the Langmuir and Hill-Langmuir model, ligand binding to the scaffold is non-saturable under this condition. (D) The prewetting model is a continuum-description of adsorption of ligands with ligand-ligand interactions on a substrate. Below the so-called prewetting concentration of ligands, ligands form a thin layer in the substrate. Above the prewetting concentration, a thick layer of ligands on the substrate is formed. Above the saturation concentration for bulk phase separation of ligands, the layer thickness does not increase anymore.

**Figure S2.**
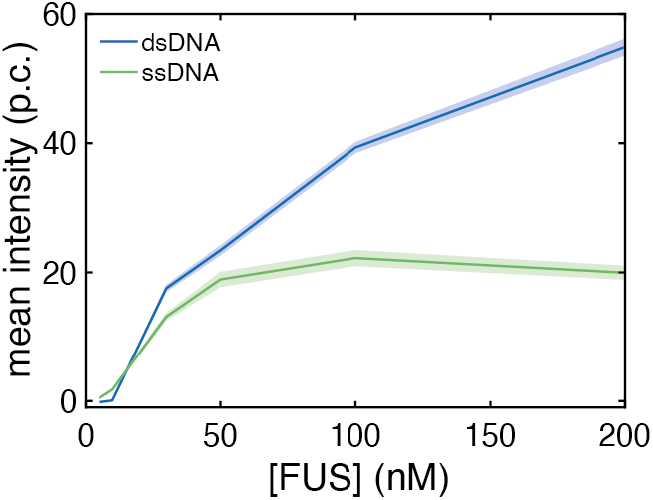
FUS adsorption on stretched DNA is saturable. Equilibrium intensity of FUS on stretched dsDNA (blue) and stretched ssDNA (green) obtained from binding experiments performed with overstretched DNA at FUS concentrations between 5 and 200 nM (p.c.: photon count; mean ± SEM)

**Figure S3.**
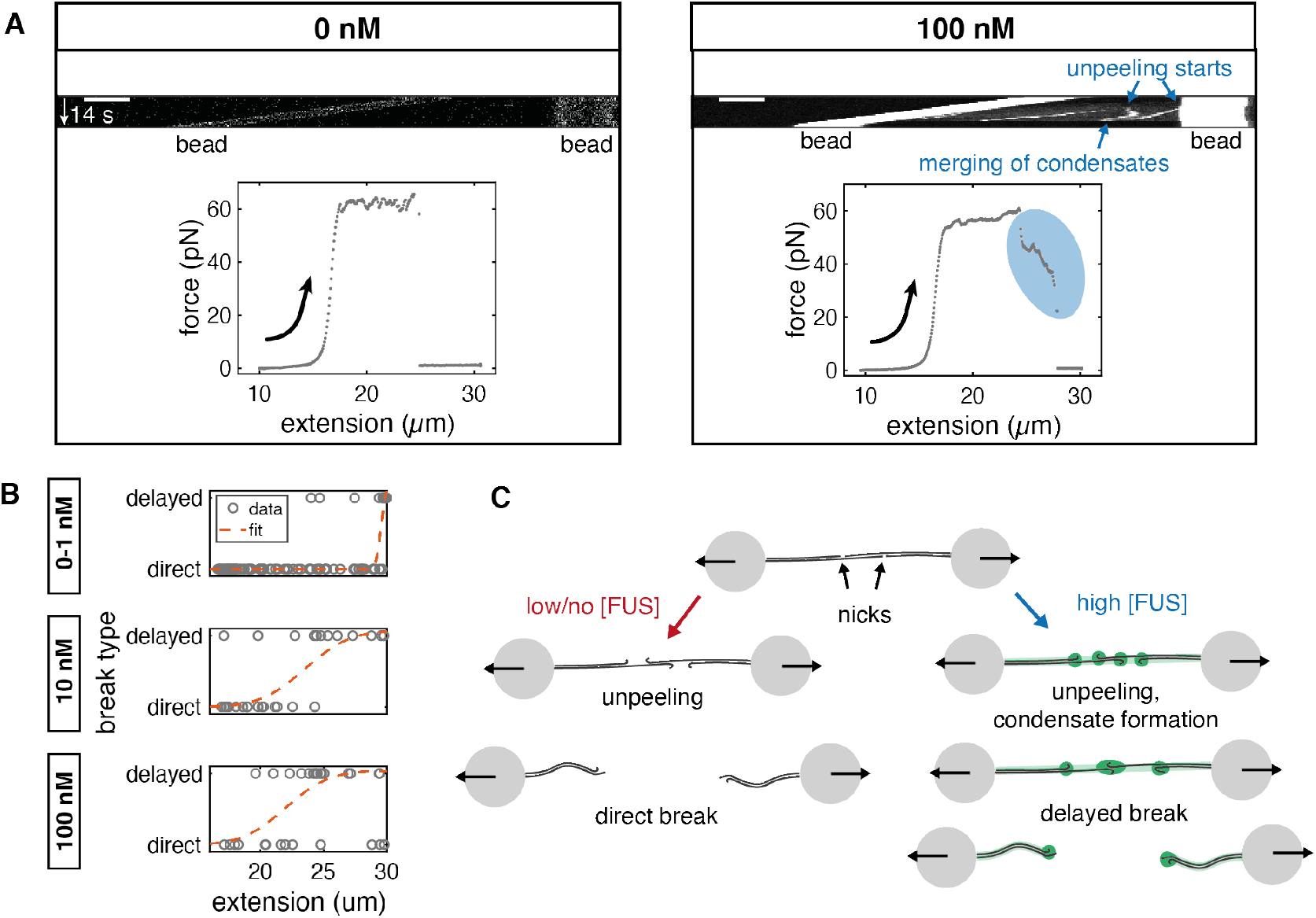
FUS adsorption delays DNA rupturing. (A) Kymographs and force-extension curves from DNA rupture experiments at 0 and 100 nM FUS. Example DNA showed a direct break in absence of FUS, while at 100 nM FUS, the break was delayed. This delay was accompanied by the fusion of two condensates moving towards each other, visible in the corresponding kymograph. Scale bar: 4 μm (B) Breaks were classified into ‘direct’ and ‘delayed’ and the DNA extension at which they occurred was measured. Error functions were fitted to estimate the characteristic extension above which delayed breaks typically occurred. Number of analyzed DNA molecules: 0/1 nM: 96, 10 nM: 29, 100 nM: 29 (C) Illustration of the rupturing process

**Figure S4.**
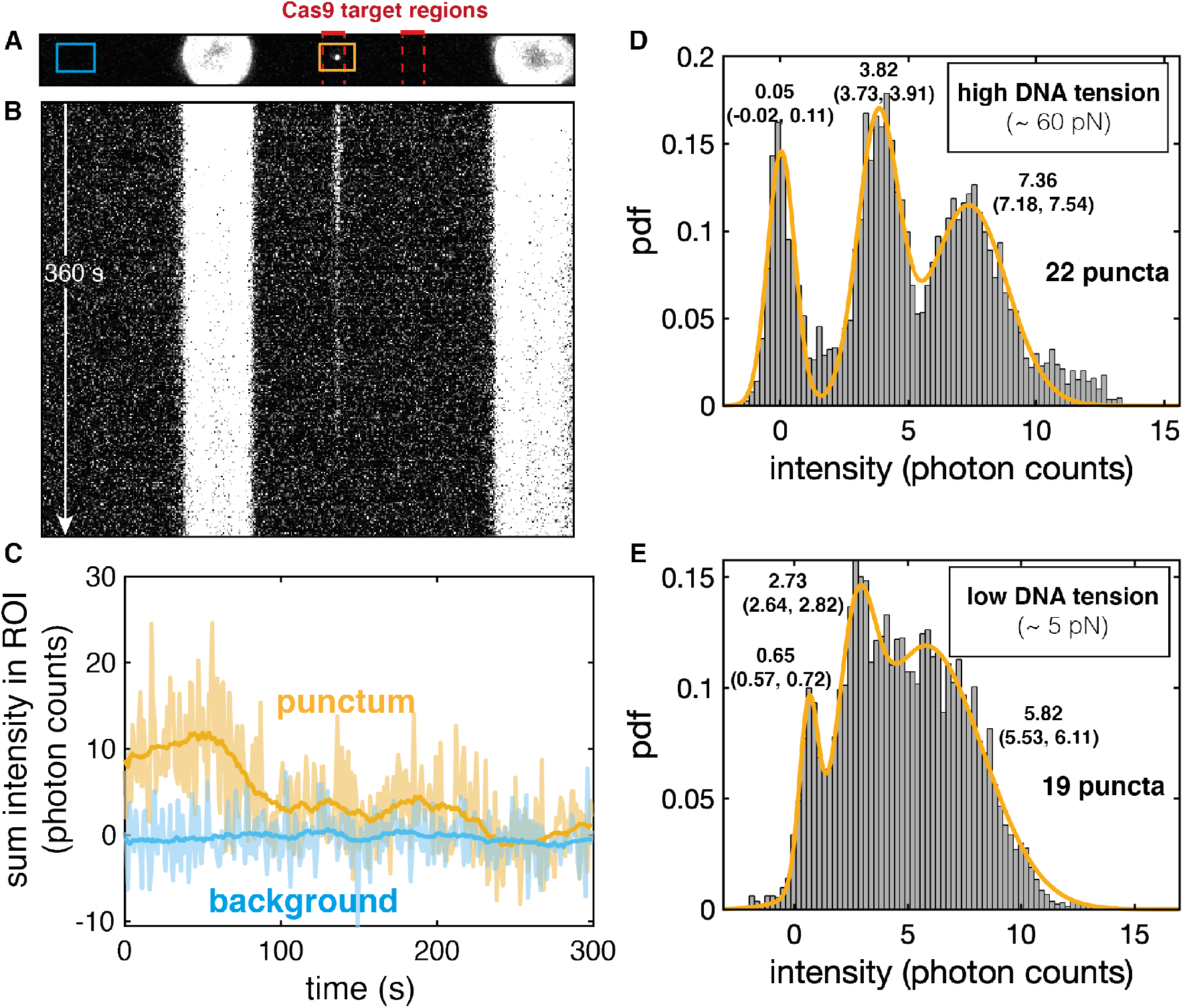
Estimation of single GFP fluorescence intensity using dCas9-GFP. (A) Representative maximum intensity projection image of dCas9-GFP binding to lambda phage DNA. dCas9-GFP was complexed with 4 different guide RNAs corresponding to 4 adjacent sequences localized at ~1/3 of the contour length of lambda phage DNA. dCas9-RNA complexes were incubated with lambda phage DNA before binding of DNA to the beads, resulting in the stable attachment of up to four complexes to the DNA target regions. Imaging was performed with the same settings as the FUS-DNA binding experiments. DNA was held either in an overstretched configuration (18 μm, ~60 pN) or in a relaxed configuration (15 μm, < 5 pN). (B) Kymograph of the experiment shown in (A), bead size: 4 μm. (C) Time traces of the summed intensity inside two segments of the imaging ROI (shown in (A). Light blue: background ROI. Orange: ROI containing a punctum that represents multiple dCas9-GFP molecules bound to adjacent sites at ~1/3 of the contour length of lambda phage DNA. The time trace of the punctum shows discrete intensity levels. Over time, intensity decreases, indicative of photo bleaching events. (Transparent lines: raw intensities, bold lines: moving average over 30 frames). (D) Histogram of intensities (moving average over 30 frames) for imaging experiments performed at high DNA tension. Three Gaussians were fitted to capture the main peaks. They represent the background intensity peak as well as the intensity of one and two GFP molecules. (22 puncta were analyzed). (E) Histogram of intensities (moving average over 30 frames) for imaging experiments performed at low DNA tension. Three Gaussians were fitted to capture the main peaks. They represent the background intensity peak as well as the intensity of one and two GFP molecules. Peaks are less distinct due to the increased fluctuations of DNA at low tension. Moreover, the intensity found for one GFP is lower (2.73 p.c. compared to 3.82) (19 puncta were analyzed).

**Figure S5.**
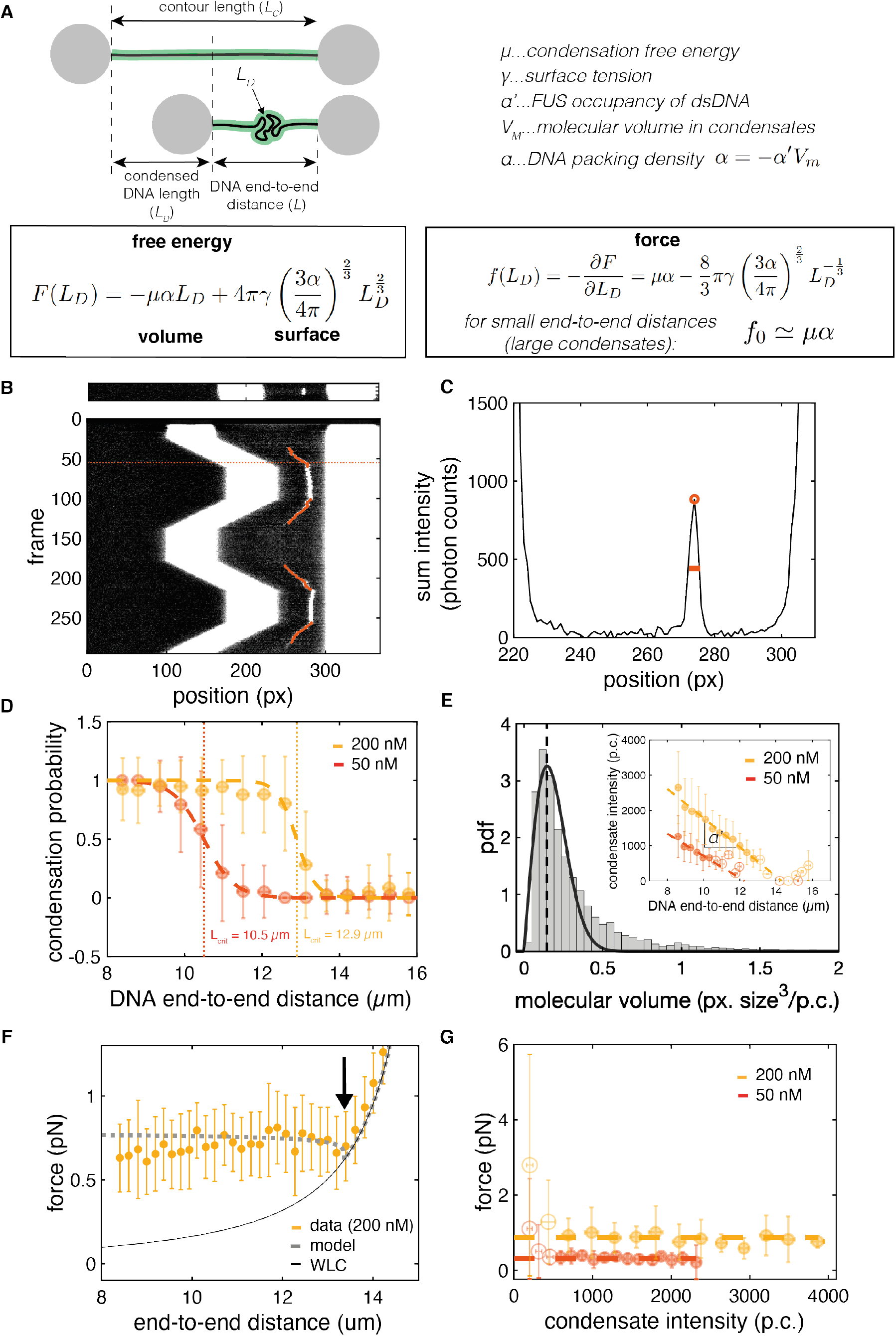
Analysis of FUS-dsDNA co-condensate formation. (A) Model for DNA condensation mediated by protein attachment. The free energy of the condensate containing the DNA length *L_D_* is determined by the volume and the surface tension of the condensate. α is the packing factor relating the condensed DNA length to the volume of the condensate. The force exerted by the condensate in order to pull in more DNA can be calculated using the negative partial derivative of the free energy with respect to *L_D_*. (B) Representative snapshot and kymograph of a FUS-dsDNA condensation experiment performed at 200 nM FUS. Overlayed in red: tracked position of the condensate when the trap position is changed. Dashed line marks the snapshot shown on top. (C) Sum intensity profile of the snapshot shown in (B). Tracked peak and FWHM are marked. For downstream analysis, we approximated the total condensate intensity as the product of peak height and FWHM. FWHM was used as an estimate of the radius of the condensate to calculate its volume (condensate approximated as sphere). (D) Probability that a condensate forms on a DNA molecule vs. DNA end-to-end distance. FUS-dsDNA condensates form below a critical, FUS concentration dependent DNA end-to-end distance *L_crit_*. Data: mean ± STD; Red: 50 nM FUS; yellow: 200 nM FUS; dashed lines: error function fits; dotted lines mark concentration dependent L_crit_. (E) Estimation of the packing factor α. α is defined as α = −α’*V_m_*. α’ is the FUS line density on DNA in FUS-dsDNA condensates. Vm is the molecular volume of FUS inside condensates. Large plot: Histogram of the probability density function (pdf) of the ratio of condensate volume and condensate intensity for all tracked condensates. Fit by a Rayleigh distribution function allows to extract the average molecular volume (volume per photon count, V_M_) by calculating the expectation value of the function. Data: histogram of 51 condensates that formed on 51 individual DNA molecules, tracked in 4109 frames; black line: Rayleigh function fitted to histogram. Inset: condensate intensity versus DNA end-to-end distance. Linear functions were fitted to the regions below *L_crit_*. Concentration dependent α’ is the slope of these functions. Data: red: condensates formed at 50 nM FUS (29 individual DNA molecules); yellow: condensates formed at 200 nM FUS (22 individual DNA molecules). Filled circles: data points classified as ‘condensate’ (below *L_crit_*); open circles: data points classified as ‘no condensate’ (above *L_crit_*). Dashed lines: linear fits indicating the linear increase of condensate size with decreasing DNA end-to-end distance. (F) Force vs. DNA end-to-end distance curve obtained from minimization of the total free energy (for details see supplementary experimental procedures). A dip at the transition from WLC to the constant force regime is observed in the experimental data and is captured by a small, but finite surface tension of the condensate of around 0.15 pN/μm (marked by black arrow). Yellow: experimental data at 200 nM FUS, mean ± STD. Dotted grey line: theory curve. Black thin line: Worm-like chain model of naked DNA (G) Force vs. condensate intensity plot. Condensates over a broad range of sizes (intensities) coexist at a constant, FUS concentration dependent force. Yellow and red: same data as shown above. Dashed lines: linear fit to the horizontal region, indicating the constant force regime.

**Figure S6.**
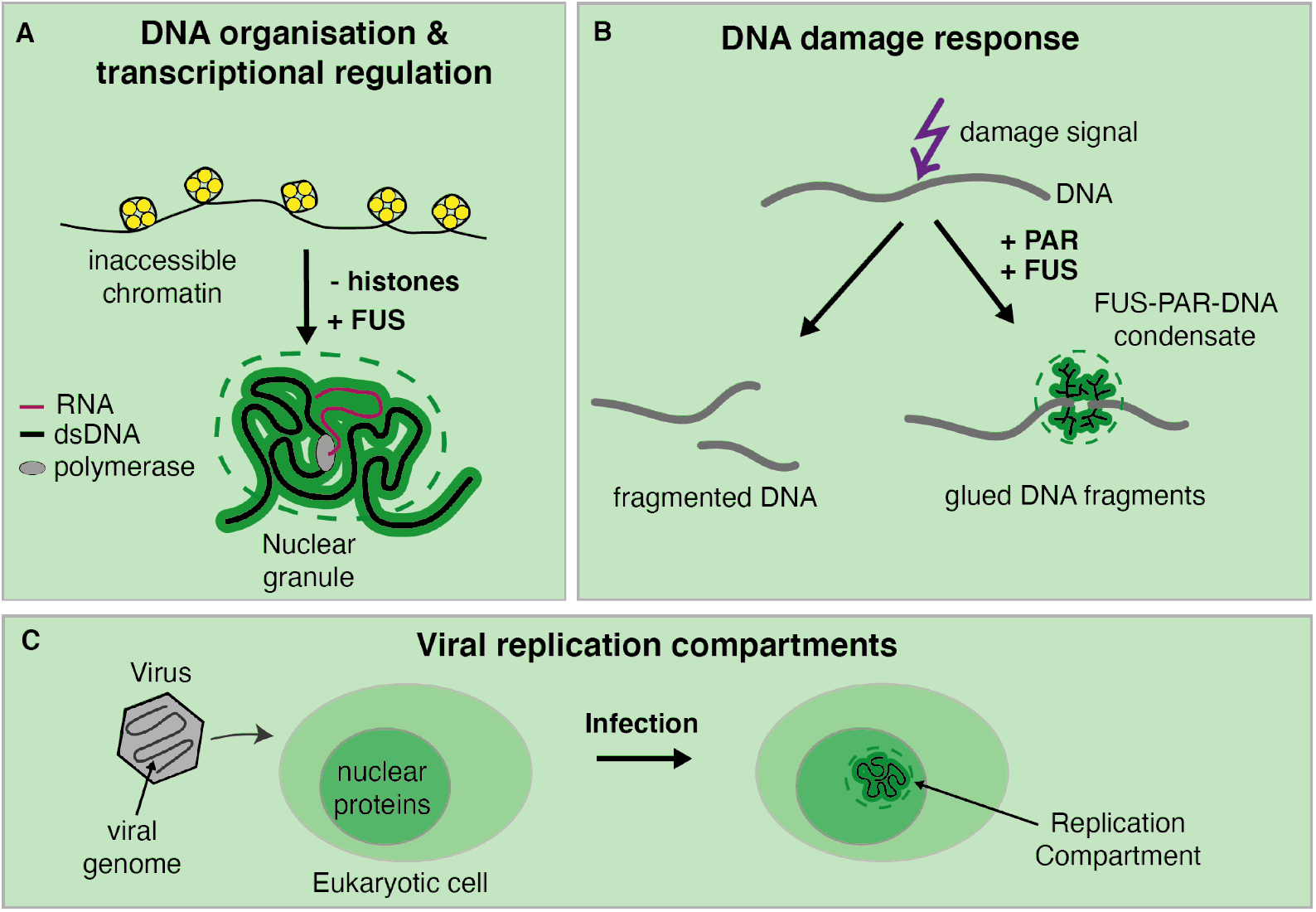
Potential physiological relevance of protein-nucleic acid co-condensation. The mechanism of monolayer protein-nucleic acid co-condensation might be the basis for the formation of (A) dynamic organizational units of highly accessible dsDNA, for example in the context of transcriptional regulation; (B) glue-like inducible DNA damage compartments at PARylated DNA damage sites, or (C) membrane-less Replication Compartments formed by highly accessible viral genomes with hijacked nuclear proteins of infected cells.

## Experimental procedures

### Protein purification

Recombinant full-length FUS-EGFP and FUS-ΔLCD-EGFP were expressed and purified as described previously (Patel et al. 2015).

### Optical tweezers experiments

Optical tweezers experiments to study the interaction of FUS-GFP (“FUS”) with DNA were performed using the fully integrated C-Trap G2 (Lumicks, Amsterdam) setup. This instrument combines optical micromanipulation via up to 4 optical traps with confocal fluorescence imaging and microfluidics flow chambers. Experimental work flows (microfluidics, trap steering, imaging settings) were controlled using the Bluelake software (Lumicks, Amsterdam). The microfluidics setup consisted of the μFlux system and a 4 or 5-channel glass chip connected via FEP tubing (1/16″ x 0.010″) (all Lumicks, Amsterdam). All experiments were carried out using custom Python scripts and at a constant temperature of 28 °C to maximize reproducibility.

Proteins were diluted in FUS buffer (70 mM KCl, 10 mM Tris, pH 7.4) to the final concentration, typically between 1 and 200 nM. Double-stranded lambda phage DNA that was biotinylated at the termini of one of the two complementary single strands (Lumicks, Amsterdam) was diluted to about 20 pg/μL in FUS buffer. 4.4 μm Streptavidin coated polystyrene beads (Spherotech) were diluted to 3 ‰ (m/v) in FUS buffer. 1 mL of each solution as well as 1 mL of plain FUS buffer were then transferred to the corresponding 4 separate channels of the μFlux system of the C-Trap. To reach a stable protein concentration in the protein channel of the flow cell we flushed the liquids at 0.8 bar for at least 45 min. Once stable experimental conditions were reached, the actual experiments were initiated.

To tether single DNA molecules, a mild flow was generated by applying a pressure of 0.2-0.3 bar. Beads were trapped in the corresponding channel and moved into the buffer channel. There, in order to estimate the stiffness of the optical traps, the thermal calibration was performed in absence of buffer flow using the in-built thermal calibration routine of the Bluelake software. The beads were then moved to the DNA channel to fish for DNA tethers. For that, in presence of mild flow, the bead-to-bead distance was periodically increased and decreased using the in-built Ping-Pong function of the Bluelake software while the force on the beads was monitored. When a characteristic force increase in response to increasing bead-to-bead distance was detected, the beads were moved back to the buffer channel. There, in absence of flow, we probed whether the tether was a single DNA molecule by measuring its force-extension curve (FEC) and comparing it with the typical FEC of lambda phage DNA. If the tether was not a single DNA molecule (or in any other way irregular), the bead pair was discarded and the routine was started again by catching a new pair of beads. If the tether was found to be a single DNA molecule, we continued with the actual experiment. DNA molecules were stretched or relaxed by changing the bead-to-bead distance. ssDNA unpeeling was induced by increasing the DNA end-to-end distance above the contour length of lambda phage DNA (~16.5 μm).

### Scanning confocal fluorescence imaging

For all fluorescence imaging experiments, the power of the 488 nm excitation laser was set to 5 % (resulting in an output of 2.14 μW) and the dwell time per pixel (pixel size 100 nm x 100 nm) to 0.05 ms. The size of the Region of Interest (ROI) was, depending on the experiment, chosen such that it could fit the DNA, the central bead segments and a region on the left side of the left bead that allowed to estimate the average background fluorescence intensity for each frame. The frame rate was set with respect to the time scales of interest in the corresponding experiment on one hand and to minimize photodamage on the other hand and thus varied between 2 and 0.25 frames per second (fps).

### Buffer exchange experiments

Individual lambda phage DNA molecules were stretched to 20 μm extension inside the buffer channel, leading to unpeeling of ssDNA starting from free ends at nicks and the DNA termini. This resulted in DNA molecules that consisted of segments of stretched dsDNA and ssDNA (at 65 pN) and relaxed ssDNA protruding from the tether at the interfaces of the stretched segments. For the binding process, the overstretched DNA molecules were then transferred into the protein channel while fluorescence imaging at the for this ROI size highest possible frame rate of 2 fps was performed for 60 s. To study the unbinding of FUS from DNA, the individual DNA molecules were transferred back to the buffer channel while imaging at 1 frame every 4 seconds (0.25 fps) was performed for 480 s in total. The reduced imaging frequency was chosen in order to minimize photo damage during these long experiments. Typically, an additional binding experiment (re-binding, same settings as initial binding) was then performed to study the reversibility of FUS-DNA interaction.

### Step-wise overstretching experiments

Individual DNA molecules were transferred into the protein channel and the bead-to-bead distance (and hence the extension of the DNA) was increased in steps of 1 μm at 5 μm/s every 10 s from initially 16 μm until the molecule broke. Imaging was performed at 1 fps.

### dsDNA flow-stretch experiments

Individual beads were held in a single trap and briefly (5-10 s) incubated in the DNA channel in presence of mild flow (0.1 bar applied) to catch individual DNA molecules attached via only one end. The beads were subsequently moved to the protein channel while the flow was maintained. During this process, imaging was performed at rates of about 1 fps.

### Repetitive dsDNA relaxation experiments

Individual DNA molecules were transferred to the protein channel. Starting from an initial extension of 16 μm, they were relaxed to 8 μm at 0.5 μm/s. After a waiting period of 20 s, the molecules were stretched to 16 μm. This was followed by another 20 s waiting period, relaxation to 8 μm and another 20 s waiting period. Finally, the bead-to-bead distance was first increased to 31 μm to rupture the molecule and then again decreased to 8 μm to estimate the force base line. During the whole experiment, fluorescence imaging was performed at 2 fps and force and bead-to-bead distance were recorded.

### dsDNA zippering experiments

Three Streptavidin coated polystyrene beads were trapped in three optical traps in a triangular configuration and moved to the DNA channel (beads 1, 2 and 3). In presence of mild buffer flow, the two beads that were aligned in parallel to the flow direction were used to fish for a DNA tether using the Ping-Pong function of the Bluelake software (beads 1 and 2). When the formation of a tether was detected using the force signal, the three beads were moved to the buffer channel again. The beads then were moved to the protein channel. Simultaneously, fluorescence imaging with a rate of approx. one frame every 5 seconds was started. The low imaging frequency was due to the large ROI required for this experiment. Once the beads were transferred to the protein channel, DNA got coated with FUS. Occasionally, an additional single DNA tether between bead 3 and 2 was formed during the process of transferring the beads from the DNA channel to the protein channel. In these cases, we straightened the tether between bead 1 and 2 by setting a bead-bead distance of around 16 μm. Further, we approached bead 3 towards the tether between bead 3 and 2 in order to enable contacting of the two FUS coated DNA tethers. We periodically approached and retracted bead 3 from the tether between beads 1 and 2 to see if potentially occurring FUS mediated zippering effects of the two DNA tethers were reversible.

### General data handling

Data analysis was performed using custom Matlab (Mathworks) routines. Image representation was performed using FIJI v. 1.51h.

For quantification of FUS intensities on DNA, background-subtracted images were generated. For that, the average background intensity of FUS in solution (obtained from regions of the image far away from beads and DNA) was computed for every frame of a time series and then subtracted from the intensity of every individual pixel of the corresponding frame.

Intensity profiles along the DNA direction were calculated by summing up background subtracted pixel intensities orthogonally to the DNA direction. Kymographs were generated by plotting the intensity profile of each frame versus the frame number.

### Analysis of buffer exchange experiments

Buffer exchange experiments were performed (1) to study the kinetics and equilibrium properties of FUS-DNA interaction and (2) to study shape changes of FUS-ssDNA condensates over time.

To study kinetics and equilibrium properties of FUS-DNA interaction, kymographs were manually segmented into regions of stretched dsDNA, stretched ssDNA and puncta (FUS-ssDNA condensates). Segmentation was done according to plausibility of the intensity pattern in terms of the DNA overstretching model and the expected relative intensity values. Intensity-time traces of each DNA segment were calculated by averaging of the intensities of all pixels in a segment for each frame. Average intensity-time traces of the different types of DNA (stretched ssDNA, stretched dsDNA, puncta) for binding and unbinding experiments performed at different FUS concentrations were calculated by averaging the intensity-time traces of every segment obtained for the corresponding experiment type (binding or unbinding) at the corresponding FUS concentration.

To extract unbinding rates, the average intensity-time traces obtained from unbinding experiments were fitted using single or double exponential functions. Fitting was performed from the time point at which the background intensity dropped (indicating that the DNA had left the protein channel) to the last time point of the experiment (480 s). The quality of fitting (represented by the R^2^ value) drastically improved by using double exponentials instead of single exponentials, particularly at elevated FUS concentrations.

Equilibrium intensities of FUS on stretched ssDNA and stretched dsDNA (i.e. the line density of FUS on DNA) were calculated by averaging the intensity-time traces obtained from binding experiments performed at different FUS concentrations over the last 30 s (i.e. when the equilibrium was reached).

To study shape changes of FUS-ssDNA condensates over time, segments of ‘puncta’ were obtained from kymographs of FUS binding experiments as described above. A custom peak finding algorithm was used to obtain the maximum intensity and the width of puncta in each frame of an experiment. The total intensity of a punctum was calculated as the product of maximum intensity and peak width.

For ensemble analysis, the individual time traces of maximum intensity, total intensity and punctum width were normalized to their final value (last 10 s of the experiment). Puncta were classified according to whether they rounded up in the course of the binding experiment. A punctum was classified as “rounded” if the normalized final maximum intensity (last 10 s of the experiment) was at least higher than the normalized initial maximum intensity (first 10 s after punctum formation) plus four times the corresponding standard deviation. Mean time traces of maximum intensity, total intensity and punctum width were calculated according to this classification.

### Analysis of FUS mediated ssDNA adhesion

Increasing the DNA end-to-end distance leads to progressive conversion of dsDNA to ssDNA via unpeeling from free ssDNA ends. Nicks in the ssDNA backbones of the dsDNA molecules define boundaries of potential ssDNA fragments. During overstretching, progressive unpeeling from the fragment boundaries will lead to dissociation of ssDNA fragments.

For analysis, we evaluated events in which two unpeeling fronts propagated towards each other when the DNA end-to-end distance was increased. When two of these fronts met and fused, they subsequently either disappeared from the field of view, indicating that the corresponding ssDNA fragment detached from the rest of the DNA molecule, or stayed attached to the rest of the tethered DNA molecule. According to this behavior, we classified events into ‘attached’ or ‘detached’. We only considered an event if the DNA tether remained intact (did not break) for at least one more step of the step-wise increase of DNA end-to-end distance.

### Analysis of FUS-ssDNA co-condensate composition

When a dsDNA molecule does not have nicks in its backbones, ssDNA unpeeling during overstretching will exclusively occur from the DNA termini, leading to the formation of exactly two FUS-ssDNA condensates on the DNA tether in presence of FUS. In this case, the number of nucleotides unpeeled from each of the two ends of the molecule and hence incorporated into each of the two condensates can be calculated from the distance between a condensate and the corresponding bead. This distance divided by the length of a ssDNA nucleotide at 65 pN (0.58 nm per nucleotide) yields the number of nucleotides in the corresponding condensate.

For every step of a step-wise overstretching experiment, in which a suitable unpeeling event occurred, we calculated the intensity profile along the DNA molecule and selected the positions of the beads and the boundaries of the condensates. From this information we calculated the integrated intensity of a condensate and the corresponding number of incorporated nucleotides. We plotted this intensity over the corresponding number of nucleotides for experiments performed in the concentration range between 1 and 200 nM FUS.

For each FUS concentration we fitted the relation between condensate intensity and number of nucleotides with a linear function. The slope of this function determines the FUS-GFP intensity per nucleotide in a condensate at a given FUS concentration and hence serves as a proxy for the ratio between protein and nucleotides in a condensate.

We plotted the slopes with respect to the corresponding FUS concentration and subsequently fitted a Langmuir isotherm in the form of 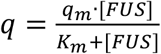 to the data (*q_m_* being the saturation occupancy of nucleotides with FUS, *K_m_* being the FUS concentration at which the occupancy has reached half of its maximum value, *[FUS]* being the FUS concentration).

### Analysis of FUS-dsDNA co-condensates

When dsDNA was relaxed in presence of sufficiently high concentrations of FUS (> 30 nM), FUS-dsDNA condensates formed. While in few instances multiple condensates formed, for analysis we focused on events where only a single condensate formed per single dsDNA molecule. We recorded image stacks for two consecutive relax-stretch cycles at 2 fps.

The free energy *F* of a FUS-dsDNA condensate containing the DNA length L_D_ can be described using a volume contribution and a surface contribution (Figure S5A), building on a framework introduced in (Quail et al. 2020):

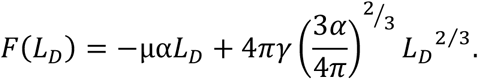

μ is the condensation free energy per volume, α is the packing factor relating L_D_ to the condensate volume, γ is the surface tension. The force required to extract a piece of DNA from the condensate is

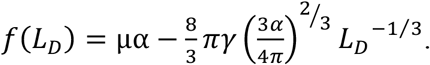

For small surface tension and high values of L_D_ (corresponding to low DNA end-to-end distances), this expression approaches a constant DNA tension *f*_0_ ≈ μα.

To analyze the mechanical properties and to finally estimate the condensation free energy per FUS molecule in FUS-dsDNA condensates, for each frame of a stack, we extracted the position, width (FWHM) and maximum intensity of the condensate using a custom peak finding algorithm (Figure S5B). The total intensity of a detected condensate in each frame was calculated as the product of maximum intensity and peak width (Figure S5C).

To correct for the base-line of the force signal, each experiment was concluded by rupturing of the DNA molecule (increase of bead-to-bead distance to 31 μm) and subsequent approach of the untethered beads to 7 μm bead-bead distance. The corresponding force signal served as a base line that was subtracted from the force signal recorded in presence of the DNA tether.

The base-line subtracted force-distance signal was synchronized with the fluorescence imaging data. For that, the raw force-distance signal (~ 9 Hz) was down sampled to 2 Hz.

The intensity of tracked condensates for every frame in which a condensate was detected was correlated to the force at which it existed and to the corresponding DNA end-to-end distance. Downstream analysis was restricted to the part of the process where the initially stretched dsDNA molecule was relaxed from 16 to 8 μm end-to-end distance (unless indicated differently). Subsequent stretching and relaxation processes in presence of FUS appeared to alter the mechanical properties of the DNA.

We first investigated at which DNA end-to-end distances FUS-dsDNA condensates exist (Figure S5D). For that we analyzed the probability to find a condensate at the different DNA end-to-end distances between 16 and 8 μm during the initial relaxation process. The step-like shapes of the curves were fitted with error functions in the shape of

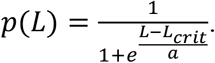

p is the probability to find a condensate, L is the DNA end-to-end distance, L_crit_ is the critical DNA end-to-end distance below which condensates typically form.

Packing factor *α* was obtained as the product of the negative slope of the linear relation between condensate intensity and DNA end-to-end distance *α*’ and the molecular volume *V_M_* of FUS in condensates (Figure S5E). *V_M_* was obtained from the expectation value of a Rayleigh distribution fit to the histogram of the ratio between volume and intensity of each detected condensate in each frame it was detected. For calculating the condensate volume, condensates were assumed to be spherical with a diameter equal to the FWHM obtained from tracking. *α’* was obtained from a linear fit to the condensate intensity vs. DNA end-to-end distance at DNA end-to-end distances below L_crit_.

The number of FUS molecules bound per DNA length inside condensates was estimated using *α’* and the intensity of a single GFP. This yielded ~115 FUS molecules bound per μm of condensed DNA at 50 nM FUS (corresponding to a spacing of one FUS molecule every ~26 bp) and ~150 FUS molecules bound per μm of condensed DNA at 200 nM FUS (corresponding to a spacing of one FUS molecule every ~20 bp).

The critical force *f_0_* was finally obtained as the mean force exerted by the condensates below L_crit_ (Figure 6H). μ as energy per volume was obtained by dividing the critical force by the corresponding packing factor for 50 and 200 nM FUS. To estimate μ of a single FUS molecules, we approximated the number of FUS molecules per μm^3^ using the molecular volume (in units of μm^3^ per photon count) and the intensity of a single GFP molecule (in units of photon counts per molecule) extracted from Figure S4. To convert into units of k_B_T per FUS molecule we assumed a temperature of 303 K and hence a conversion relation of 1 k_B_T = 4.2e-3 pNμm.

### Estimation of number of FUS molecules per condensate

To estimate the number of FUS-EGFP molecules in a condensate (Figure S4) we calibrated the fluorescence intensity using tightly bound Cas9-EGFP molecules in DNA as introduced in Morin *et al.* 2020. In brief, Cas9-EGFP was incubated with 4 different types of guide RNA molecules that had sequences complementary to 4 adjacent sequences at about 1/3 of the length of lambda phage DNA. The formed complexes were incubated with lambda phage DNA molecules so that up to 4 Cas9-EGFP molecules could tightly attach to the DNA molecules. Individual pre-incubated lambda DNA molecules were imaged at the conditions used for FUS-DNA experiments for 360 s (pixel size 100 nm, pixel dwell-time 0.05 ms, frame rate 1 fps). DNA molecules were held either at ~60 pN or at below 5 pN to allow for fluorescence calibration that could either be used for FUS-ssDNA condensates (formed on top of overstretched DNA tethers) or for FUS-dsDNA condensates (present at forces below 5 pN). The time traces of background subtracted sum intensities of puncta found in the DNA target regions were extracted (moving average over 30 frames). Multiple Gaussian distributions were fit to the probability density function (pdf) of intensities. The position of the first peak (after the ‘background peak’) was used as an estimate of the intensity of a single EGFP at either low or high DNA tension.

## Supplementary experimental procedures

### Analysis of the influence of FUS on DNA rupturing behavior

To study whether FUS influences the rupturing behavior of DNA, individual DNA molecules were transferred into the protein channel. Their extension was continuously increased at 1 μm/s until they broke, starting from 10 μm (Figure S3). Imaging was performed at 1 fps. Breaks observed in the force-extension curves were classified according to the extension at which they occurred and whether they occurred directly (force drop from overstretching plateau to zero within few data points) or in a delayed manner (via multiple intermediate states). The type of breaking event vs. the extension at which it occurred was the plotted for the FUS concentration range between 0 and 100 nM. A characteristic extension for the switch from direct to delayed breaks was estimated using an error function fit. At 100 nM a group of data points indicating direct breaks at around 30 μm extension was excluded from the fit as they probably were associated with rupturing of the DNA at the junctions with the beads rather than being caused by DNA unpeeling.

### dCas9-EGFP preparation, imaging and intensity analysis

Recombinantly expressed dCas9-EGFP was stored at 5.3 mg/ml at −80°C in storage buffer (250 mM HEPES pH 7.3, 250 mM KCl) and thawed 1 h prior to the experiment. sgRNAs were made using an in vitro expression kit against the following four adjacent target loci on lambda DNA + NGG PAM sequence GGGAGTATCGGCAGCGCCAT TGG, GGAGGATTTACGGGAACCGG CGG, GGCAACCAGCCGGATTGGCG TGG, GGCGGTTATGTCGGTACACC GGG. The spacing between adjacent target sequences was adjusted to 40 to 50 bp to prevent steric hindering of adjacent dCas9-sgRNA complexes. The target region marked by the 4 adjacent RNA sequences corresponds to a region at 1/3 (or 2/3) of the DNA contour. Guide RNAs were expressed and purified using commercial kits (MEGAscript T7 Transcription Kit, Invitrogen and mirVana miRNA isolation Kit, Invitrogen); stored in ddH_2_O at 0.6 – 1 mg/mL at −80°C and thawed together with the dCas9-EGFP protein 1 h prior to the experiment.

First, 2 μL of dCas9-EGFP were pre-diluted into 38 uL complex buffer (20mM Tris-HCl pH 7.5, 200 mM KCl, 5 mM MgCl_2_, 1 mM DTT) prior to the complexing reaction. Second, 5 μL of the 20x dCas9-EGFP dilution were mixed with 4 μL sgRNA stock which contained all four sgRNAs in equal stoichiometries. Subsequently, the reaction volume was adjusted to 50 μL by adding 41 μL complex buffer and incubated at room temperature (22°C) for 30 min.

After incubation was completed, the 10 μL of the dCas9-sgRNA complex reaction are mixed with 1 μL of 5 nM biotinylated lambda DNA. The reaction volume was then adjusted to 50 μL by adding 39 μL reaction buffer (40 mM Tris-HCl pH 7.5, 200 mM KCl, 1 mg/mL BSA, 1 mM MgCl_2_ and 1 mM DTT) followed by a second incubation for 30 min at room temperature (22°C).

Lambda phage DNA molecules were diluted in FUS buffer and transferred to the microfluidics system of the C-Trap setup. Individual DNA molecules equipped with dCas9-guide RNA complexes were tethered as described before. Fluorescence imaging was performed at 1 fps over 360 frames with a pixel size of 100 nm, a pixel dwell time of 0.05 ms and 5% intensity of the 488 nm excitation laser. DNA molecules were held at a tension of either ~5 pN (‘low tension’) or ~60 pN (‘high tension’).

For image analysis, every frame was background subtracted. Fluorescent puncta sitting at 1/3 or 2/3 of the DNA contour length were segmented and the total intensity within these ROIs was extracted for each frame of each experiment (Figure S4). Note that puncta that were observed outside of the DNA target regions at 1/3 or 2/3 of the DNA contour length were not considered for analysis as they were suspected to represent dysfunctional and hence potentially aggregated dCas9-EGFP.

For DNA held at low or high tension, the probability density function (pdf) of the intensity inside the ROI in each frame was represented in a histogram. Notably, only up to 3 or 4 clear peaks were visible, indicating that typically not all 4 different dCas-RNA complexes were bound to the target region of the DNA. The peaks were fitted using Gaussian functions and the position of the second peak (the first one depicts the background intensity) was considered to be the approximate intensity of a single EGFP molecule under the corresponding imaging conditions (at high DNA tension: 3.82 ± 0.09 p.c.; at low DNA tension: 2.72 ± 0.09 p.c. (95% confidence interval)).

### Towards a full model of the FUS-dsDNA co-condensates held in optical traps

To capture the full force vs. end-to-end distance curve of the FUS-dsDNA condensate in a dual trap optical tweezer experiment, we consider a free energy that contains contributions from the FUS-DNA condensate, the stretched DNA polymer, and the optical traps. The free energy *F_cond_* of the condensate is written as

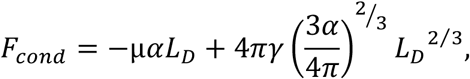

where μ denotes the condensation free energy per volume, L_D_ is the length of DNA contained in the condensate, α is the packing factor relating L_D_ to the condensate volume, and γ is the surface tension (Quail *et al.*, 2020). The mechanical energy F_WLC_ stored in a stretched DNA polymer is determined by integration of the polymer force f_WLC_ (Worm Like Chain, WLC) (Smith, Cui and Bustamante, 1996) according to

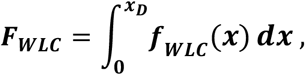

with

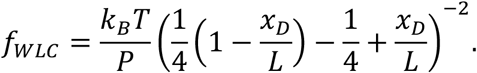

Here, T denotes the absolute temperature, k_B_ is Boltzmann’s constant, L is the contour length of the DNA that is not condensed (L = L_C_ – L_D_ with L_C_ denoting the contour length of 16.5 μm of the entire piece of lambda phage DNA), and x_D_ is the DNA end-to-end distance and thus the separation distance between the surfaces of the two beads in the optical trap. The energy stored in the two optical traps is given by

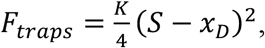

where K denotes the stiffness of the optical traps and S is the distance between the two trap centers, minus twice the radius of the beads. Hence, S – x_D_ denotes the summed displacement of the two beads in the two optical traps. The total free energy of the system F is now given by

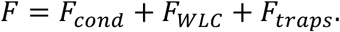

We consider the ensemble where the distance S between the two trap centers is fixed, and x_D_ and L_D_ are fluctuating quantities that approach values that correspond to a minimum of F. We determine this minimum numerically for the following values of the input parameters: L_C_ = 16.5 μm, k_B_T = 4.15 pNnm, P = 50 nm, μ = 11.78 pN/μm^2^, α = 0.059 μm^2^, γ = 0.15 pN/μm, K = 0.15 pN/nm. For a pair of values of L_D_ and x_D_ at which F is minimal, we determine the DNA tension according to 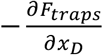 or partial 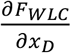. The resulting force vs. end-to-end distance curve was plotted on top of the experimental data obtained for 200 nM FUS (Figure S5F).

### Supplementary movies

Movie S1

Continuous overstretching of lambda phage DNA in presence of 100 nM FUS showing homogeneous adhesion of FUS to stretched ssDNA and dsDNA and formation of condensates of FUS with unpeeled ssDNA (bead size: 4 μm)

Movie S2

Repetitive overstretching of lambda phage DNA in presence of 100 nM FUS, showing reversibility of FUS-ssDNA condensate formation (bead size: 4 μm)

Movie S3

Step-wise overstretching of lambda phage DNA in presence of 100 nM FUS, indicating viscoelastic-like material properties of FUS-ssDNA condensates (bead size: 4 μm)

Movie S4

A single lambda phage DNA molecule attached to a bead and stretched by hydrodynamic flow. Attachment of FUS (100 nM) leads to DNA condensation (bead size: 4 μm)

Movie S5

Repetitive stretch-relax cycles of lambda phage DNA in presence of 100 nM FUS at end-to-end distances below the DNA contour length. Reversible formation of a FUS-dsDNA condensate is observed (bead size: 4 μm)

Movie S6

FUS-mediated reversible zippering of two dsDNA strands studied using three optical traps (bead size: 4 μm)

## Notes

### Competing Interest Statement

The authors have declared no competing interest.

